# Photoinactivation of catalase sensitizes *Candida albicans* and *Candida auris* to ROS-producing agents and immune cells

**DOI:** 10.1101/2021.08.31.458449

**Authors:** Pu-Ting Dong, Yuewei Zhan, Sebastian Jusuf, Jie Hui, Zeina Dagher, Michael K. Mansour, Ji-Xin Cheng

## Abstract

Nearly all organisms found in nature have evolved and developed their own specific strategies to cope with reactive oxygen species (ROS). Catalase, a heme-containing tetramer protein expressed in a broad range of aerobic fungi, has been utilized as an essential enzymatic ROS detoxifying mechanism, and shows remarkable efficiency in degrading hydrogen peroxide (H_2_O_2_) for fungal cell survival and host invasion. Here, we demonstrate that catalase inactivation with blue light renders fungal cells highly susceptible to ROS attack, thus resembling a ‘strength-to-weakness optical switch’. To unveil catalase as the underlying molecular target of blue light and its inactivation mechanism, we systematically compared wild-type *Candida albicans* to a catalase-deficient mutant strain for susceptibility to ROS in the absence/presence of 410 nm treatment. Upon testing on a wide range of fungal species and strains, we found that intracellular catalase could be effectively and universally inactivated by 410 nm blue light. We find that the photoinactivation of catalase in combination with ROS-generating agents is highly effective and potent in achieving full eradication of multiple fungal species and strains, including multiple clinical strains of *Candida auris*, the causative agent of the global fungal epidemic. In addition, photoinactivation of catalase is shown to facilitate macrophage killing of intracellular *Candida albicans*. The antifungal efficacy of catalase photoinactivation is further validated using a *Candida albicans*-induced mouse model of skin abrasion. Taken together, our findings offer a novel catalase-photoinactivation approach to address multidrug-resistant *Candida* infections.

## Introduction

The frequency of invasive fungal infections in immunocompromised patients has been consistently increasing over the past few decades (1). *Candida* species are the most common cause of human fungal infections (2), such as oropharyngeal, cutaneous candidiasis and mucosal or deep-seated organ infections (3). Invasive *Candida* infections remain a significant cause of morbidity and mortality among immunocompromised patients, partly due to the growing spread of antifungal resistance (4). The growing emergence of multidrug-resistant *Candida* species poses an alarming trend worldwide (5). Especially of concern is *Candida auris* (*C. auris*) which has resulted in multiple worldwide healthcare outbreaks with record-breaking mortality rate due to its multi-drug or even pan-drug resistance nature and biofilm formation (6, 7). Ongoing COVID-19 pandemic has further accelerated *C. auris* outbreaks (8). Life-threatening invasive mycosis has been recorded to accompany COVID-19 patients due to impaired immune responses, forcing higher than normal usage of antifungal agents (9). At a higher level, the long-term imprudent usage of antifungal agents has been shown to accelerate the resistance of anti-fungal development (need a reference here). Confronted with this severe situation, novel therapeutic approaches are highly desired.

Antimicrobial blue light, especially in the 400-430 nm range, has drawn increasing attention in recent years as a non-drug approach to treat wide-ranging bacterial (10, 11) and fungal infections (12, 13). The antifungal efficacy of blue light has been studied by many research groups worldwide. Blue light at 415 nm has demonstrated the capability to inactivate *C. albicans* both *in vitro* and *in vivo* mouse burn infection models, and the susceptibility of *C. albicans* to blue light-induced inactivation did not change even after the 10^th^ passage in the presence of blue light exposure suggesting the unlikelihood of fungi developing photoresistance (14). *Gupta et al*. achieved 4.52-log_10_ reduction of *C. albicans* after delivering 332.1 J/cm^2^ of 405 nm irradiance (15). *Rosa et al*. reported a 2.3-log_10_ reduction of *C. albicans* in a biofilm setting under the treatment of 455 nm exposure with a dose of 45.16 J/cm^2^ (16). *Raquel et al*. further demonstrated significant antimicrobial effect of 405 nm on *C. albicans* both in monomicrobial and polymicrobial biofilm schemes (17). Blue light was also utilized to combine with other agents to eliminate *C. albicans. Leon at al*. showed that quinine chloride could enhance the fungicidal effects of blue light 405 nm on the elimination of *C. albicans* both *in vitro* and *in vivo*. Besides *C. albicans*, blue light can also inhibit the growth of microconidia of molds, such as *Trichophyton rubrum* and *Trichophyton mentagrophytes* (18). Studies also demonstrated that 405 nm light has detrimental effect on additional yeasts such as *Sacchromyces cerevisiae* and obligate molds including *Aspergillus niger* (19). Noteworthy, no significant loss of viability on human keratinocytes was observed after 216 J/cm^2^ blue light treatment (17). Collectively, these studies demonstrate that blue light works effectively against major pathogenic fungal species without developing resistance and no significant cytotoxicity on host cells, thus emerging as a drug-free alternative approach to combat fungal infections.

However, the underlying antimicrobial mechanisms of blue light has stayed elusive for decades. A widely believed hypothesis is that metal-free porphyrin (in the 405-420 nm window) or riboflavin (in the 450-470 nm window) are endogenous molecular targets (12). Bacterial/fungal inactivation are eradicated by the reactive oxygen species (ROS) produced through photodynamic reaction between these molecules and blue light (20). However, this perspective remains controversial. First, it has been reported that the total amount of coprotoporphyrins inside the microbes was not a contributing factor to the antimicrobial efficacy of blue light (21). Second, the intracellular concentration of porphyrins or riboflavin is as low as 2-4× 10^−3^ mg/ml (22). Given the situation that research of antimicrobial blue light on fungi stays understudied and the working mechanism of antimicrobial blue light has yet to be clarified, other possible molecular targets are assumed to exist.

Here, using the wild type *C. albicans* SC5314 along with a catalase-deficient *C. albicans* mutant, we demonstrate that catalase is a major molecular target of antimicrobial blue light (400-420 nm window). Catalase from wide-ranging fungal species could be effectively inactivated by 410 nm blue light. Subsequently, photoinactivation of catalase renders fungal cells highly susceptible to non-fungicidal low-concentration hydrogen peroxide (H_2_O_2_) and ROS-producing antifungal drugs. Moreover, the synergy between photoinactivation of catalase and H_2_O_2_ can enable total eradication of multiple notorious drug-resistant *C. auris* isolates whereas either alone has limited efficacy. Furthermore, photoinactivation of catalase substantially boosts macrophages activity against *C. albicans* indicated by shorter hyphae length and higher eradication percent of *C. albicans* were observed in the case of 410 nm involved group. Finally, we validated the synergistic effect between photoinactivation of catalase and H_2_O_2_ in a *C. albicans*-induced mouse skin abrasion model without noticeable skin damage. Collectively, our findings provide a novel catalase-targeting strategy to treat multi-drug resistant fungal infections.

## Results

### Catalase inside fungal cells can be inactivated by blue light irradiance

As shown in **Figure 1a**, catalase (from bovine liver) revealed by PyMOL has a tetramer structure, with porphyrin rings hiding inside its hydrophobic pocket. It was reported as early as in 1965 that bovine liver catalase can be inactivated by visible light exposure (23). Nonetheless, photoinactivation of catalase remained underexplored over the decades. To better understand how photons inactivate catalase, especially catalase inside fungal cells, we first quantified the remaining catalase percentage from bovine liver catalase (2.5 U/ml) after blue light exposure under various wavelengths using an Amplex red catalase kit. And we found that blue light, especially in the 400-420 nm window, inactivates catalase by 70% compared to control (**Figure 1b**). 410 nm shows the best capability to inactivate catalase, which might be due to the fact that the optimal absorption peak of catalase is around 410 nm. In the following experiments, we used continuous-wave (CW) 410 nm LED as the light source to inactivate catalase.

**Figure 1.**
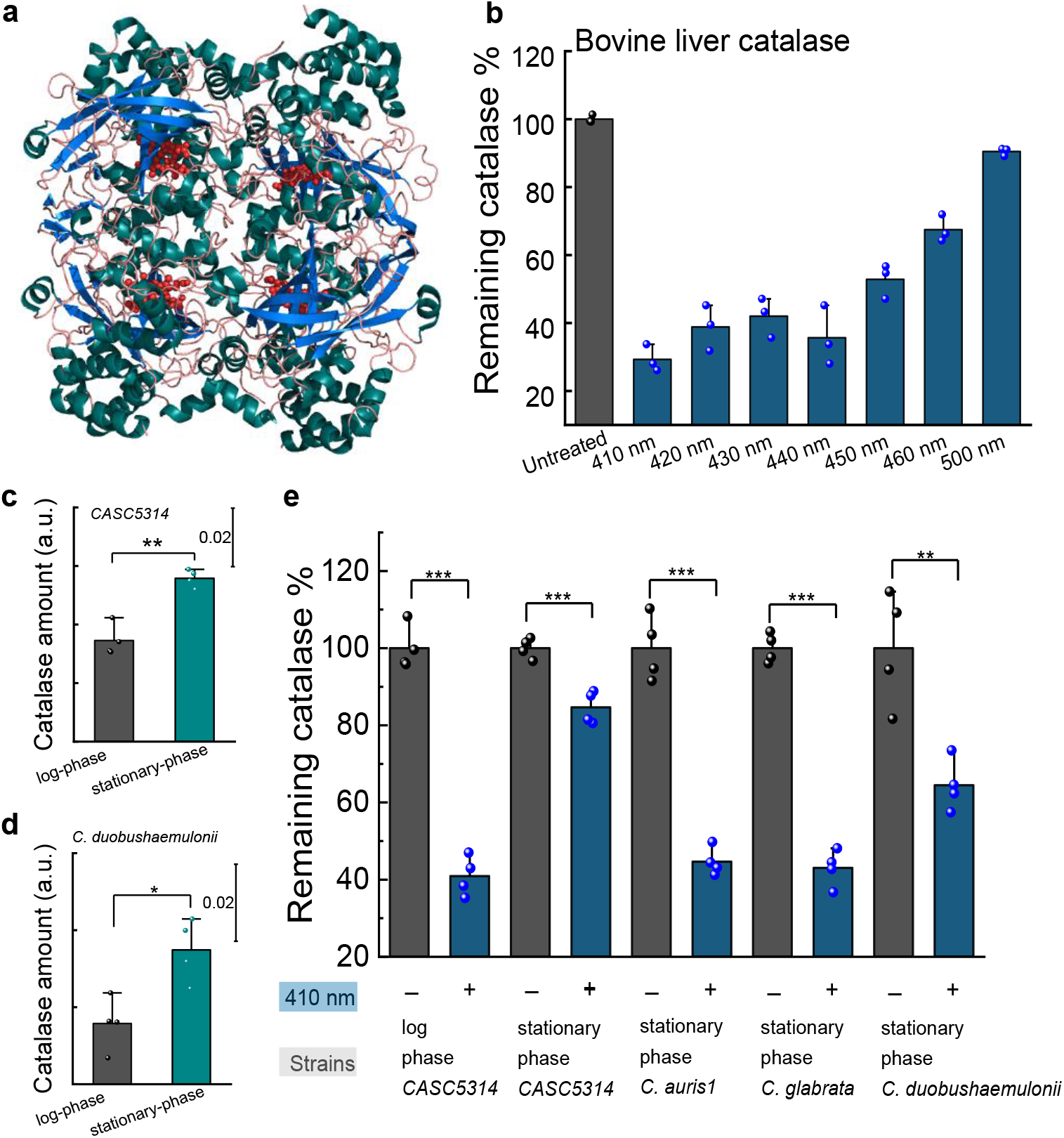
Catalase from bovine liver and fungi cells can be inactivated by blue light. **a**. Structure of bovine liver catalase revealed by PyMOL simulation. PDB ID: 1TGU. **b**. Quantification of remaining catalase percentage of bovine liver catalase (2.5 U/ml) by Amplex Red catalase assay under different treatment schemes. N=3. Dose: 15 J/cm^2^. **c-d**. Comparison of catalase amount between log- and stationary-phase *C. albicans* SC5314 (**c**) and *Candida duobushaemulonii* (**d**). **e**. Quantification of remaining catalase percentage in various fungal strains with and without 410 nm blue light. Dose: 30 J/cm^2^. Data: Mean+SD from at least three replicates. Student unpaired *t*-test. *: *p*<0.05, **: *p*<0.01; ***: *p*<0.001.

To query whether fungal catalase can be inactivated by blue light in the same way as liver catalase, we tested the remaining catalase percentage inside the fungal cells after 410 nm exposure. Prior to that, we investigated the catalase expression level for fungal cells at different metabolic phases. It was reported that stationary-phase microbes usually have higher amount of sigma factor *σ^S^*, the key protein responsible for survival and improved resistance under stressful conditions (24). Increased expression of *σ^S^* leads to the relatively higher amount of catalase in the stationary-phase microbes (24). As shown in **Figure 1c-d**, we indeed found that stationary-phase *Candida spp*. have more catalase compared to log-phase. Of note, catalase in log- and stationary-phase *C. albicans* could be consistently inactivated by 410 nm exposure. Moreover, this inactivation behavior exists in many other fungal strains apart from *C. albicans* (**Figure 1e**). In short, we found that catalase inside fungal cells can be effectively inactivated by 410 nm exposure regardless catalase expression level.

### Photoinactivation of catalase sensitizes *C. albicans* to H_2_O_2_

Having established that fungal catalase can be efficiently inactivated by 410 nm blue light, we then queried whether this attenuation effect sensitizes *C. albicans* to H_2_O_2_ considering the major function of catalase. A colony-forming unit (CFU/ml) assay was conducted by combining 410 nm with H_2_O_2_ at a non-fungicidal concentration (22 mM). As shown in **Figure 2a**, photoinactivation of catalase significantly sensitizes wild-type *C. albicans* SC5314 to H_2_O_2_ by around six orders of magnitude, whereas H_2_O_2_ or blue light alone didn’t have apparent killing effect. This result indicates a synergistic killing effect between photoinactivation of catalase and H_2_O_2_. To understand whether this augmented killing was due to intracellular accumulation of H_2_O_2_ after catalase inactivation, we utilized an established kit using H_2_O_2_-sensitive fluorophore to quantify intracellular H_2_O_2_ in *C. albicans* cells treated with H_2_O_2_ or 410 nm plus H_2_O_2_ through a confocal laser scanning microscope (CLSM). As shown in **Figure 2b-c**, fluorescent intensity in the 410 nm plus H_2_O_2_ treated single *C. albicans* SC5314 was significantly higher than cells treated by H_2_O_2_ alone, suggesting that photoinactivation of catalase boosts intracellular H_2_O_2_ accumulation.

**Figure 2.**
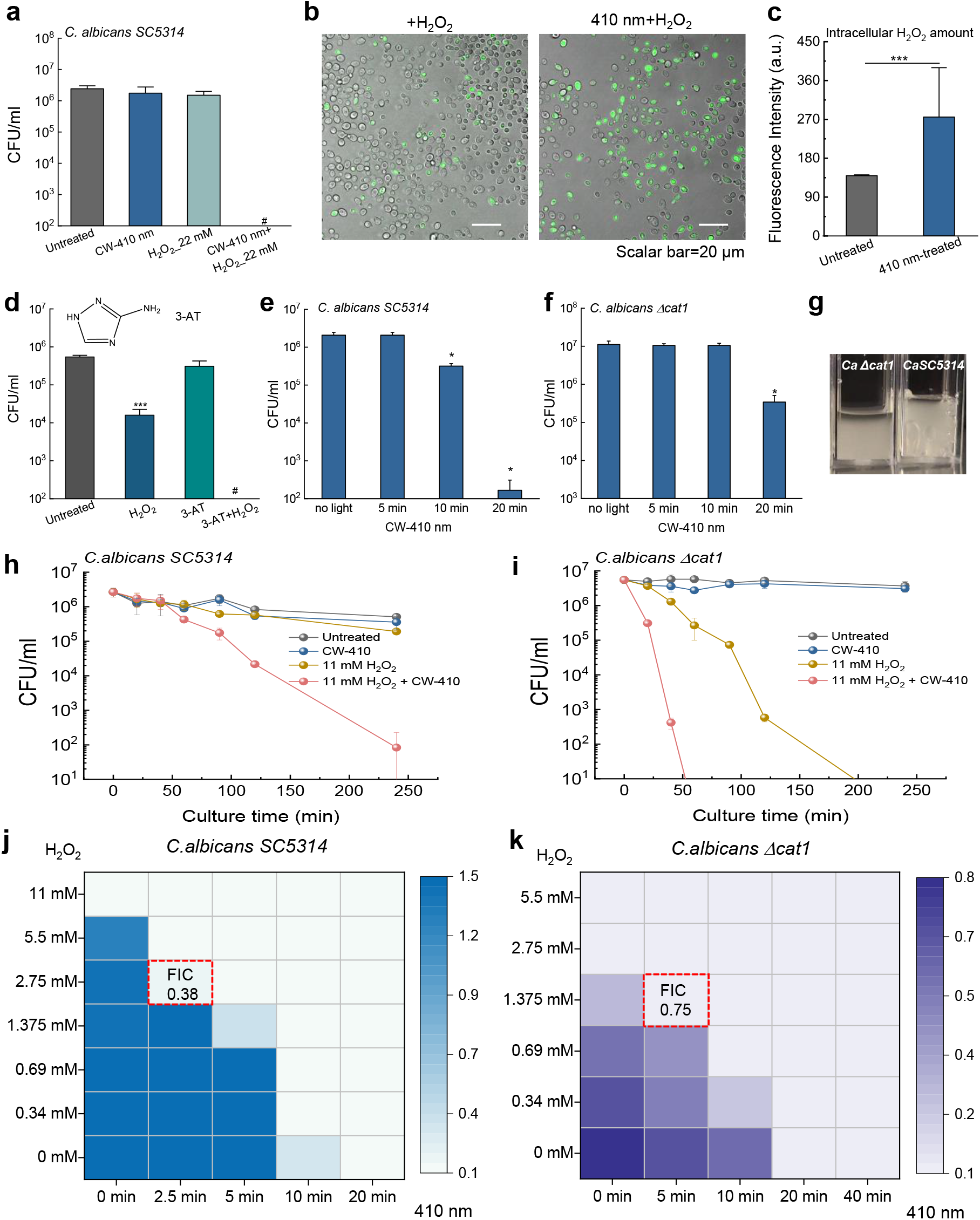
Photoinactivation of catalase sensitizes *C. albicans* to H_2_O_2_ and catalase is identified as a primary molecular target of blue light. **a**. Colony-forming unit (CFU/ml) of *C. albicans* SC5314 under different treatment schemes. **b**. Confocal imaging of intracellular H_2_O_2_ with/without 410 nm exposure. Scalar bar=20 μm. H_2_O_2_: 22 mM, images were taken after 30-min incubation and washing step. **c**. Quantitation of intracellular H_2_O_2_ via fluorescence intensity from images acquired in **b**. **d**. CFU/ml of *C. albicans* SC5314 in the presence of 3-AT. Significant difference from the untreated group. 3-AT: 50 mM, 4-hour incubation. **e-f**. CFU/ml of *C. albicans* SC5314 and its catalase-deficient mutant *C. albicans Δcat1* under 410 nm exposure with various doses. **g**. Pictures showing bubble formation phenomena of *C. albicans Δcat1* and *C. albicans SC5314* in the presence of 3% H_2_O_2_. **h-i**. Time-killing curves of *C. albicans* SC5314 (**h**) and *C. albicans Δcat1* (**i**) under different treatment schemes. **j-k**. Checkerboard broth dilution assay showing the combinatorial behavior between blue light and H_2_O_2_ against *C. albicans* SC5314 and *C. albicans Δcat1*. Data: Mean+SD. N=3. Student unpaired *t*-test. *: *p*<0.05, **: *p*<0.01; ***: *p*<0.001. Pound sign (#) means CFU results are below the detection limit.

To investigate whether catalase is the primary target for blue light exposure, we conducted the following experiments. First, a catalase inhibitor, 3-Amino-1,2,4-triazole (3-AT) (25), was utilized to chemically inhibit catalase activity within *C. albicans*, and then the susceptibility to H_2_O_2_ was recorded. As shown in **Figure 2d**, total eradication was obtained when adding H_2_O_2_ to 3-AT treated *C. albicans* SC5314 whereas 3-AT alone only exerts limited killing efficiency, suggesting the pivotal role that catalase plays in neutralizing H_2_O_2_. Second, we exploited a catalase-deficient *C. albicans* strain, *C. albicans Δcat1*, to further evaluate the function of catalase underlying the antimicrobial effect of blue light. *C. albicans Δcat1* demonstrates significantly lower susceptibility to 410 nm blue light compared to its the wild type (**Figure 2e-f**), indicating catalase-mediated sensitivity to 410 nm blue light.

Next we examined the time-kill CFU assay of wild type *C. albicans* SC5314 and catalase-deficient *C. albicans Δcat1* under different treatment schemes. *C. albicans Δcat 1* did not produce prototypical oxygen bubbles as the wild type *C. albicans SC5314* did in the presence of H_2_O_2_ (**Figure 2g**), confirming the major role of catalase in H_2_O_2_ neutralization. As shown in **Figure 2h**, four hours after treatments, both H_2_O_2_ and 410 nm blue light alone didn’t exert significant fungicidal effects, whereas H_2_O_2_ plus 410 nm blue light-treated group had around a 5-log_10_ reduction of wild-type *C. albicans* SC5314. These data suggest that there is an ample synergy between photoinactivation of catalase and H_2_O_2_ against *C. albicans*. Of note, when applying the same H_2_O_2_ with concentration used to test wild type *C. albicans* SC5314, H_2_O_2_ alone achieved more than 5-log_10_ reduction of *C. albicans Δcat1* (**Figure 2i, supplementary Figure 1a**). Moreover, *C. albicans Δcat1* exhibited similar susceptibility to H_2_O_2_ killing compared to wild type *C. albicans* SC5314 exposed to 410 nm plus H_2_O_2_ in a time-kill assay, corroborating that catalase is a key target of 410 nm light. Noteworthy, enhanced H_2_O_2_ killing of *C. albicans Δcat1* after 410 nm exposure was observed (**Figure 2i, supplementary Figure 1b**), hinting those additional molecular targets other than catalase might exist and be responsive to 410 nm exposure.

To further confirm the synergistic effect between photoinactivation of catalase and H_2_O_2_, we conducted a checkerboard broth dilution assay to derive a fractional inhibitory concentration index (FICI) (26). FICI was calculated through the following equation:

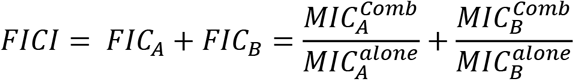
where 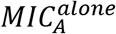 and 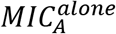 are the minimal inhibitory concentrations (MICs) of the drugs A and B when functioning alone, and 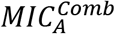 and 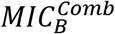 are the MICs of drugs A and B in combination, respectively. A synergy between two agents was defined with FICI ≤ 0.5 (27), and ‘no interaction’ with a FICI in the range of 0.5-4 (27). The MICs of 410 nm against *C. albicans* SC5314 and *C. albicans Δcat1* were 282 J/cm^2^ and 564 J/cm^2^, respectively (**supplementary Figure 2a, c**). And MICs of H_2_O_2_ were 11 mM and 2.75 mM, respectively (**supplementary Figure 2b, d**). Once combining 410 nm and H_2_O_2_ together, MIC of H_2_O_2_ against wild type *C. albicans* SC5314 was around 2.75 mM (**Figure 2j**), and 1.375 mM against *C. albicans Δcat 1* (**Figure 2k**). FICI from both strains could be calculated from the above equation. Then we obtained a FICI of 0.38 in the case of wild type *C. albicans* SC5314, and a FICI of 0.75 for *C. albicans Δcat1*. These results indicate that there is an effective synergy between photoinactivation of catalase and H_2_O_2_ in eradicating *C. albicans*.

### Photoinactivation of catalase enhances ROS-generating agents to inhibit the proliferation of wide-ranging fungal cells

After demonstrating the synergy between photoinactivation of catalase with non-fungicidal low-concentration H_2_O_2_ against *C. albicans*, we next query whether this synergy still holds effective against other fungal species. To have a high-throughput analysis, we adopted a PrestoBlue viability assay (28) to investigate the time-course proliferation of wide-ranging fungal species under different treatment schemes (**supplementary Figures 3-4**).

As shown in **Figure 3a**, photoinactivation of catalase consistently enhances the fungistatic effect of H_2_O_2_ against both log- and stationary-phase *C. albicans* SC5314, and *C. albicans* C13, *C. albicans* C14 as indicated by nearly 0% proliferation rate in the combination-treated group. Besides *C. albicans*, we also investigated whether this enhancement works for non-albicans *Candida* species. *Candida glabrata* (*C. glabrata*), the commensal saprophyte, caused increasing incidence and prevalence of mucosal and systemic infections due to the widespread use of immunosuppressive and antimycotic therapy (29). *Candida parapsilosis* (*C. parapsilosis*), the second most commonly isolated *Candida* species from blood cultures, could also cause invasive *C. parapsilosis* infections in both AIDS patients and surgical patients (30). *Candida lusitaniae* (*C. lusitaniae*), fungal pathogen capable of developing both acute and long-term infections, can rapidly develop resistance (31) to multiple antifungals and lead to the breakouts of multidrug-resistant fungal infections (32). *Candida haemulonii* (*C. haemulonii*), the opportunistic fungal pathogen associated with bloodstream infections, has emerged as a multi-drug resistant yeast for deep-seated soft tissue and bone infections in diabetic patients (33). Thus, we further wondered whether photoinactivation of catalase enhances the antifungal efficacy of H_2_O_2_ against these fungal species.

**Figure 3.**
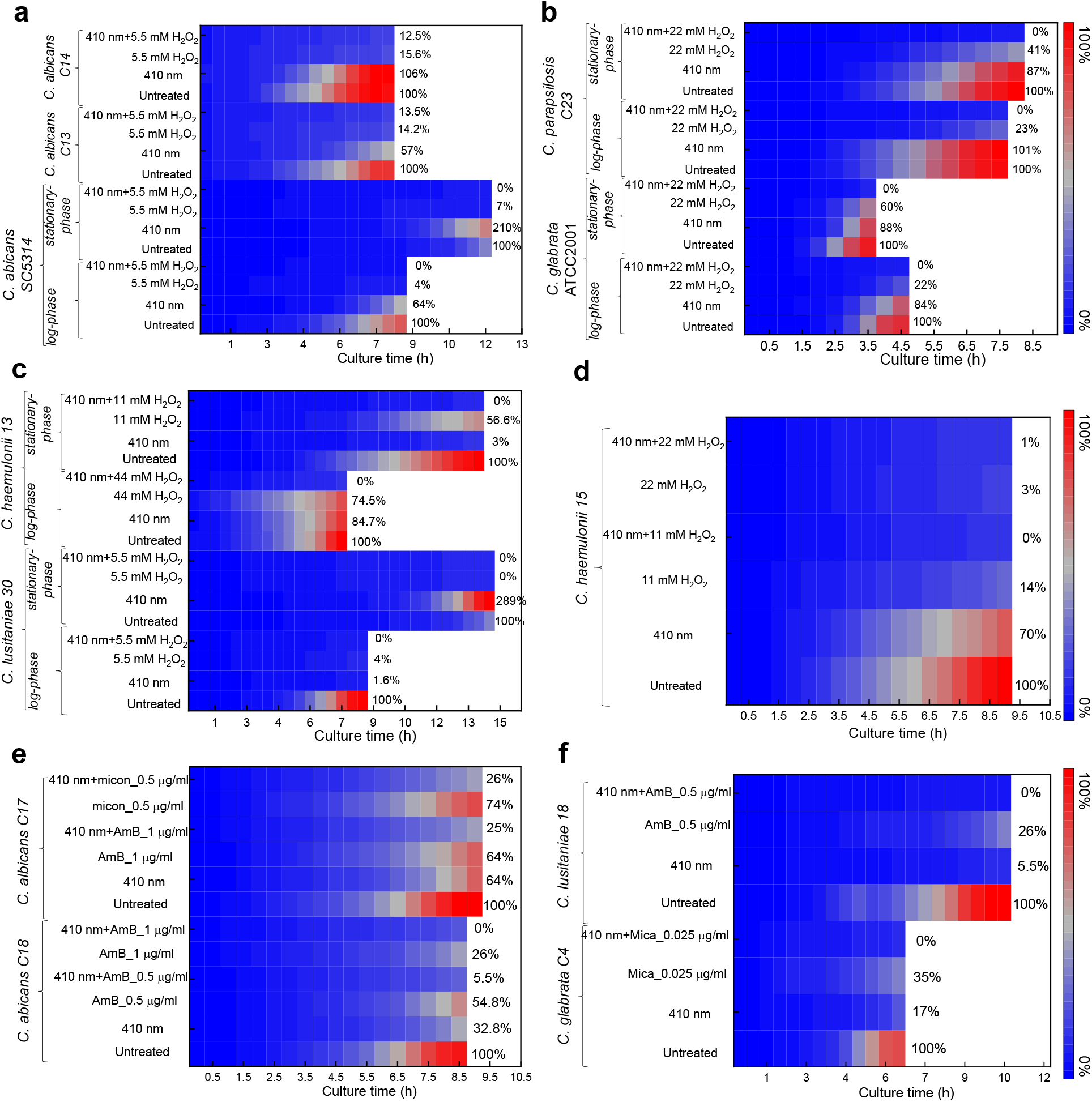
Photoinactivation of catalase enhances ROS-generating agents to inhibit the proliferation of wide-ranging fungal cells. Data were acquired through PrestoBlue proliferation assay and were presented as mean from three replicates. Blue light: 410 nm, 30 J/cm^2^. Abbreviations: AmB (amphotericin B), micon (miconazole), mica (micafungin).

Noteworthy, complete inhibition of proliferation (0% growth compared to the untreated group) was obtained from all the above mentioned fungal species in the 410 nm plus H_2_O_2_ treated group (**Figure 3, b-d; supplementary Figure 3**), both in the form of log- and stationary-phase. For example, in the case of stationary-phase *C. glabrata* ATCC2001 (**Figure 3b**), 410 nm exposure (30 J/cm^2^) inhibited around 12% of the original fungal cells to proliferate. At a concentration of 22 mM, H_2_O_2_ alone prohibited the growth of around 40% of the original fungal cells. Once combining these two treatments together, total growth inhibition was obtained. These findings consolidated the synergy between photoinactivation of catalase and H_2_O_2_ against those catalase-positive fungal pathogens, which holds clinical potential for treating multidrug-resistant fungal infections.

To treat fungal infections in the clinic, currently there are limited classes of antifungals available, such as liposomal formulations of amphotericin B (representative of polyenes), miconazole (azole representative), echinocandin (antifungal which inhibits the cell wall synthesis). Interestingly, it was reported that all these antifungals are able to induce an intracellular ROS burst within fungal cells as part of their proposed antifungal mechanism (34–36). Considering the pivotal role of catalase in scavenging ROS (37) and the fact that fluconazole and amphotericin-B resistance are associated with increased expression of catalase and superoxide dismutase (38), we reasoned that photoinactivation of catalase could enhance the antifungal efficacy of these antimycotics against fungal cells. The same time-course PrestoBlue proliferation assay was applied for the evaluation when treating fungal cells with these antifungals at a sublethal concentration (**supplementary Figure 4**).

As shown in **Figure 3e**, in the case of *C. albicans* C17, 410 nm blue light alone (30 J/cm^2^) inhibited around 36% of original *C. albicans* to divide, amphotericin B (AmB, 1 μg/ml) alone also gave out around 36% inhibition effects, whereas the administration of AmB after 410 nm blue light treatment drastically suppressed fungal growth by 75%. Similar phenomenon was found in the combinational behavior between 410 nm treatment and miconazole (micon, 0.5 μg/ml) against *C. albicans* (**Figure 3e**). Micafungin (mica, 0.025 μg/ml) has demonstrated the augmented inhibition effect induced by photoinactivation of catalase against *C. glabrata* as well (**Figure 3f**). In combination with the time-course calibration curves with known number of fungal cells under the same conditions, we can derive the relatively proliferated fungal cell numbers (**supplementary Figure 5–6**). Through comparison between the proliferated fungal cell numbers under different treatment schemes, we can clearly observe the enhanced inhibition effect by 410 nm (**supplementary Figure 6**). Collectively, these data were in line with our hypothesis, that is, photoinactivation of catalase boosts the antimycotic efficacy of commonly used antifungal agents.

### Elimination of *Candida auris* by synergizing photoinactivation of catalase with H_2_O_2_

*Candida auris* (*C. auris*), as an emerging and particularly notable *Candida* species, has been associated with nosocomial outbreaks on five continents since 2009 due to its innate resistance to multiple classes of antifungal drugs (39, 40). Its high morbidity and mortality have led to it being one of the major public health concerns worldwide. *C. auris* demonstrates a unique propensity to colonize and persist on various surfaces, contributing to multiple outbreaks in health care settings (41). *C. auris* has also been reported to cause bloodstream infections and urinary tract infections for patients with COVID-19 (8, 41), thus likely as a compounding factor in COVID-19 pandemic. Most of the clinical isolated *C. auris* strains exhibit resistance to triazoles (fluconazole) and other antifungal drugs (42). Confronted with this situation, it is imperative to have alternative approaches to both sterilize clinical settings and further treat *C. auris*-caused infections.

*C. auris* strains have been reported to be extremely resistant to hydrogen peroxide vapor sterilization (43), this triggered us to wonder whether this persistence is due to catalase. To investigate whether photoinactivation of catalase and H_2_O_2_ could efficiently kill *C. auris*, we conducted CFU assay of multiple *C. auris* clinical isolates acquired from antimicrobial resistance bank (AR-BANK) of the Centers for Disease Control and Prevention (CDC). Here *C. auris* 1, 2, 5, 6 represent AR-BANK#0381, 0382, 0385, 0386, respectively.

Log-phase *C. auris* isolates were exposed to 410 nm blue light (36 J/cm^2^) followed by subsequent administration of H_2_O_2_ (4-h incubation at 30°C, 11-22 mM), serial dilution and CFU enumeration. As shown in **Figure 4a**, in the case of *C. auris 1*, photoinactivation of catalase renders this pathogen highly susceptible to H_2_O_2_ by around three orders of magnitude. Drastically, total eradication of *C. auris 2* was obtained in the 410 nm plus H_2_O_2_ treated group (**Figure 4b**). Similar augmentation effect appeared with regard to *C. auris 5* and *C. auris 6* as well (**Figure 4c-d**). These results underline the essential role of catalase for *C. auris* to defend external H_2_O_2_ attack. Furthermore, photoinactivation of catalase works synergistically with H_2_O_2_ against clinical *C. auris*.

**Figure 4.**
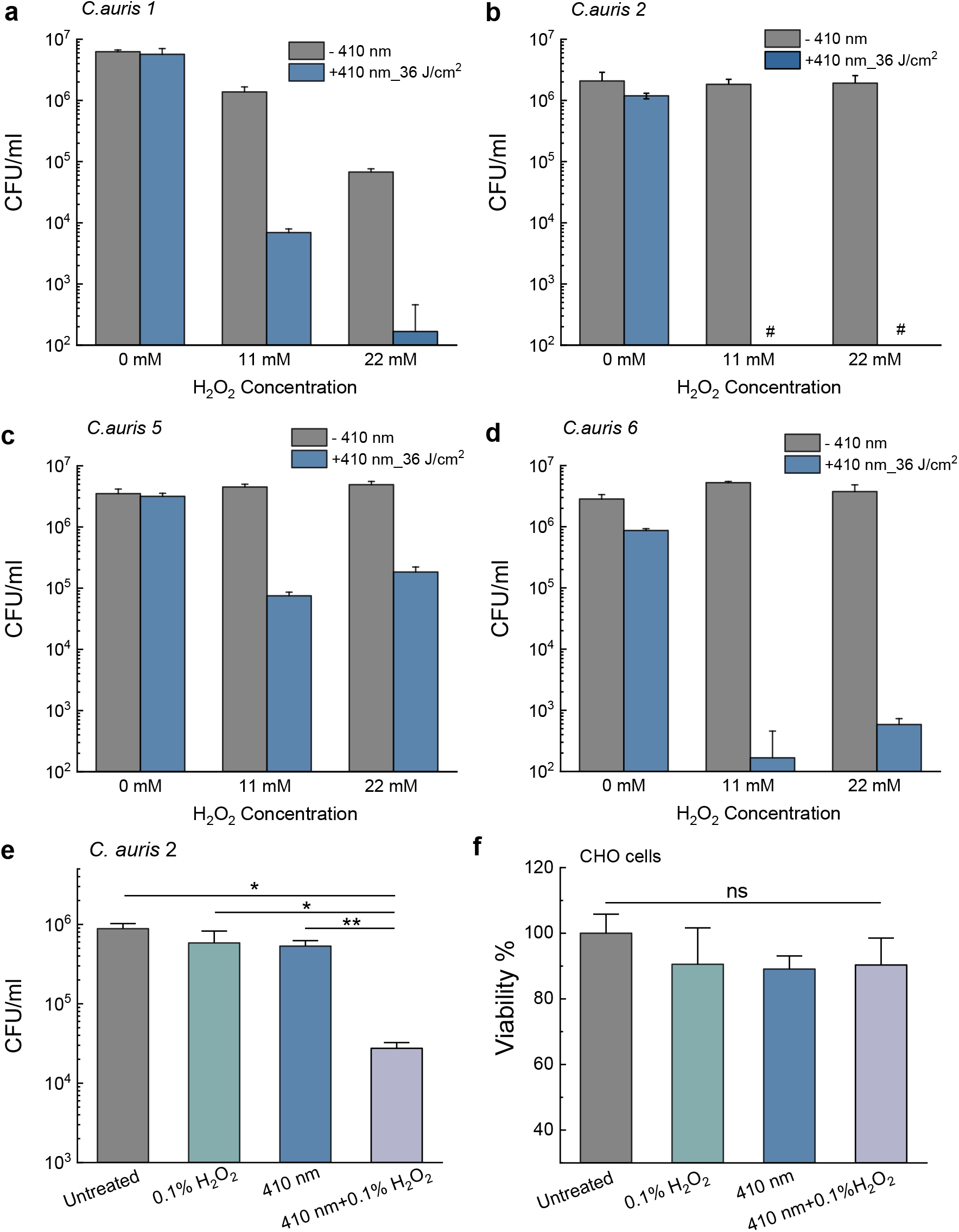
Elimination of various clinical *Candida auris* strains via synergy between photoinactivation of catalase with H_2_O_2_. CFU/ml of log-phase *C. auris 1* (**a**), *C. auris 2* (**b**), *C. auris 5* (**c**), and *C. auris 6* (**d**) under different treatment schemes. Data: Mean+SD. N=3. Pound sign (#) means below the detection limit. H_2_O_2_-incubation time: 4 hours. 410 nm: 36 J/cm^2^. (**e**). CFU/ml of *C. auris 2* under 410 nm blue light followed by short-time incubation with 0.1% H_2_O_2_ (30 mM, 1-min incubation). 410 nm blue light dose: 30 J/cm^2^. (**f**). Viability of CHO cells by MTT assay under the same treatment conditions as panel (**e**). Statistical analysis was achieved by student unpaired *t*-test. **: *p*<0.01; *: *p*<0.05; ns: not significant.

It was reported that hydrogen peroxide vapor was utilized as clinical room disinfection in order to get rid of *C. auris* colonization (44), however the *C. auris* biofilm is difficult to eradicate explaining the concerning epidemiology of high relapse rates in clinical settings (43). To query the clinical potential of this synergistic therapy which requires shortened H_2_O_2_ treatment time, we then reduced both the H_2_O_2_ incubation time and 410 nm blue light treatment time and tested if such synergistic treatment remains effective against *C. auris*. We treated 410 nm blue light-exposed *C. auris 2* with H_2_O_2_ at a higher concentration (30 mM, 0.1%) for only 1 minute, then followed by CFU enumeration. As shown in **Figure 4e**, there was no significant difference between the untreated group and either blue light alone-treated group or H_2_O_2_ alone-treated group. However, around 99% of fungal burden (**Figure 4e**) was reduced once combining these two treatments together, suggesting the consistent and effective synergy between photoinactivation of catalase and H_2_O_2_. Noteworthy, the same treatment schemes did not exert detrimental effects to a Chinese Hamster Ovary (CHO) cell line according to MTT assay (**Figure 4f**). Collectively, photoinactivation of catalase and H_2_O_2_ exhibit synergistic fungicidal effects against multiple clinical *C. auris* strains.

### Proliferation inhibition of wide-ranging *C. auris* strains by combining photoinactivation of catalase with ROS-producing agents

Due to the prominent significance of *C. auris*-caused breakouts, we particularly investigated whether photoinactivation of catalase could sensitize wide-ranging *C. auris* strains to multiple ROS-producing agents (H_2_O_2_, amphotericin B, fluconazole) in a high-throughput approach, PrestoBlue proliferation assay was utilized again to investigate the fungistatic effects under various treatment schemes (**supplementary Figure 7–11**). Meanwhile, time-course PrestoBlue fluorescence intensity of untreated fungal cells with known CFU number were recorded as standard curves. In this way, we could derive the relatively proliferated fungal cell number from the treated groups based on the linear calibration curves (**supplementary Figure 12–13**).

As shown in **Figure 5a-b**, 410 nm blue light alone (30 J/cm^2^) inhibits around the proliferation of *C. auris 1* by 90%, and blue light alone (30 J/cm^2^) significantly suppressed the growth of *Candida auris*. Total inhibition effect was obtained in the 410 nm plus H_2_O_2_ (5.5 mM) treated group. Meanwhile, we also observed that total inhibition of proliferation happened in the 410 nm plus amphotericin B (AmB) treated group whereas AmB alone exerted limited efficacy. Noteworthy, photoinactivation of catalase could enhance the fungistatic effects of fluconazole by around one order of magnitude (**Figure 5a**). Similar phenomena were observed in the case of *C. auris 3* (MIC of fluconazole is 128 μg/ml) and *C. auris 10* (MIC of fluconazole is >256 μg/ml) as well (**Figure 5c-d**). Strikingly, we further found that multiple *C. auris* strains are highly susceptible to 410 nm blue light treatment alone (**Figure 5e-f**) as total inhibition was observed, hinting that ample blue light-sensitive endogenous chromophores existing inside *C. auris* strains. Collectively, *C. auris* were highly sensitive towards blue light, and photoinactivation of catalase renders *C. auris* susceptible to various ROS-producing agents, even for some antifungal agents to which *C. auris* have developed resistance.

**Figure 5.**
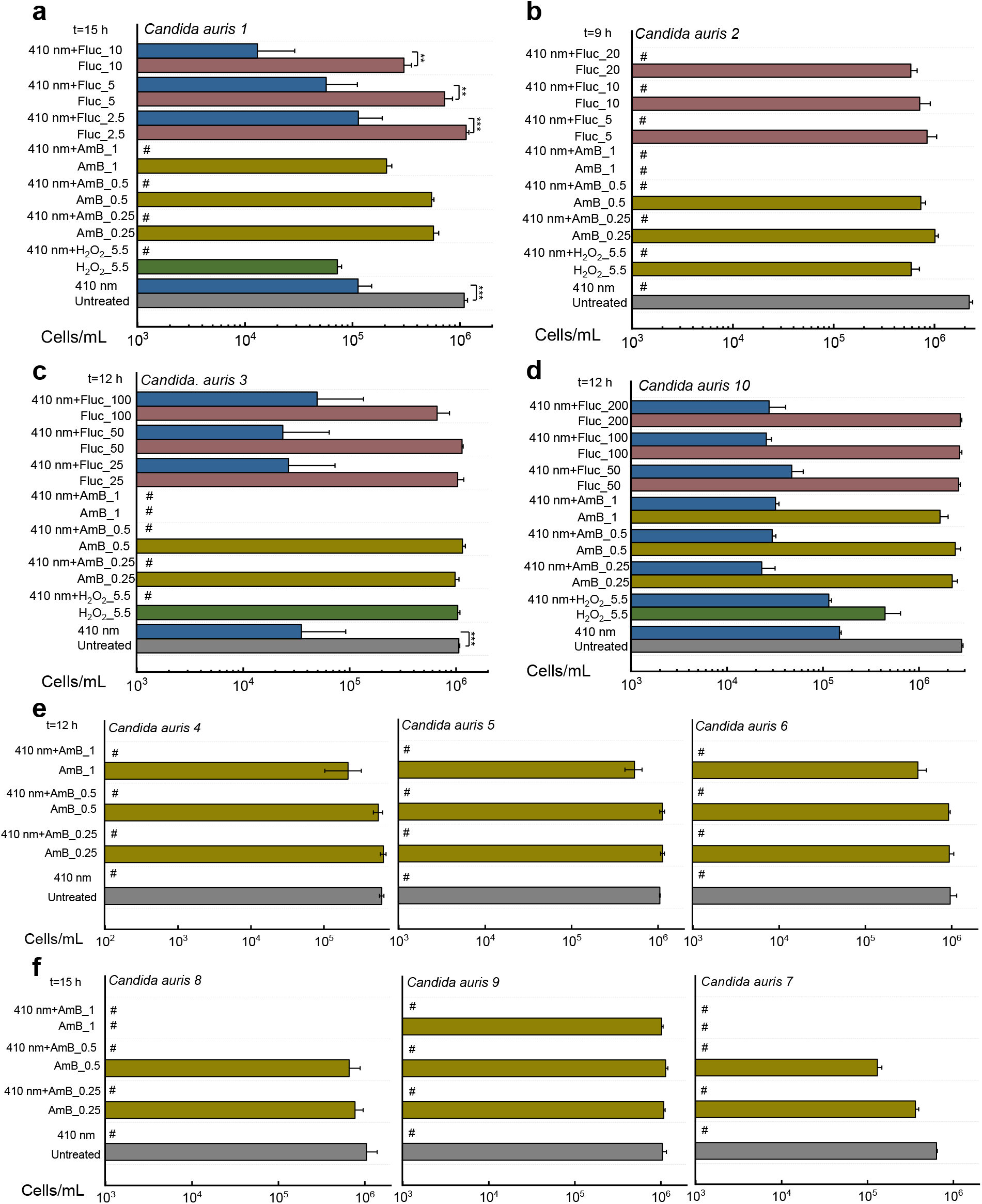
Derived cells/ml from ten clinical *Candida auris* strains based on the PrestoBlue proliferation assay and calibration curves. Data: Mean+SD. N=3. Abbreviations: amphotericin B (AmB), fluconazole (Fluc). Significant different was determined through student unpaired *t*-test. *: *p*<0.05, **: *p*<0.01; ***: *p*<0.001. Pound sign (#) means calculated values are below the detection limit.

### Photoinactivation of catalase boosts macrophage killing of *C. albicans*

When fungal infection occurs, pathogenic fungi will encounter our critical line of defense, the innate immune system where phagocytic macrophages can recognize, engulf, and destroy these fungal cells (45). It was reported that *C. albicans* can cause macrophage membrane rupture and lysis through hyphal germination (46). Intracellular *C. albicans* can survive and duplicate within human macrophages by harnessing array of strategies. One of such essential strategies is the expression of ROS scavenging enzymes, catalase and superoxide dismutase (45). Thus, we wondered whether photoinactivation of catalase could deprive *C. albicans* off this vital virulence factor, thus facilitating macrophage killing.

To test this hypothesis, we infected a macrophage cell line RAW 264.7 with untreated *C. albicans* and 410 nm blue light-treated *C. albicans* at a multiplicity of infection of 10 for 1 hour. After that, we performed a Live (SYTO 9)/Dead (PI) staining assay to visualize intracellular *C. albicans*. As shown in **Figure 6a-c**, after one hour of infection, *C. albicans* indeed exhibited hyphal morphology, and pierced through the macrophages. A small portion of *C. albicans* remained alive inside the macrophages (**Figure 6a**). By comparison, in the 410 nm blue light-treated group, we not only observed significantly less *C. albicans* in their hyphal form, but also significantly shortened hyphal length (**Figure 6d-f**). Quantitative analysis of the hyphal length under different treatment groups (**Figure 6g**) further consolidated this finding. Furthermore, there is a significant reduction of *C. albicans* viability after macrophage phagocytosed 410 nm pre-exposed *C. albicans* (**Figure 6h**). Collectively, these evidences suggest that photoinactivation of catalase attenuates the virulence of *C. albicans*, thus boosting macrophage elimination of intracellular *C. albicans*. Of note, blue light (410 nm) treatment did not pose significant toxicity to CHO cells (**Figure 6i**) under the same dosage.

**Figure 6.**
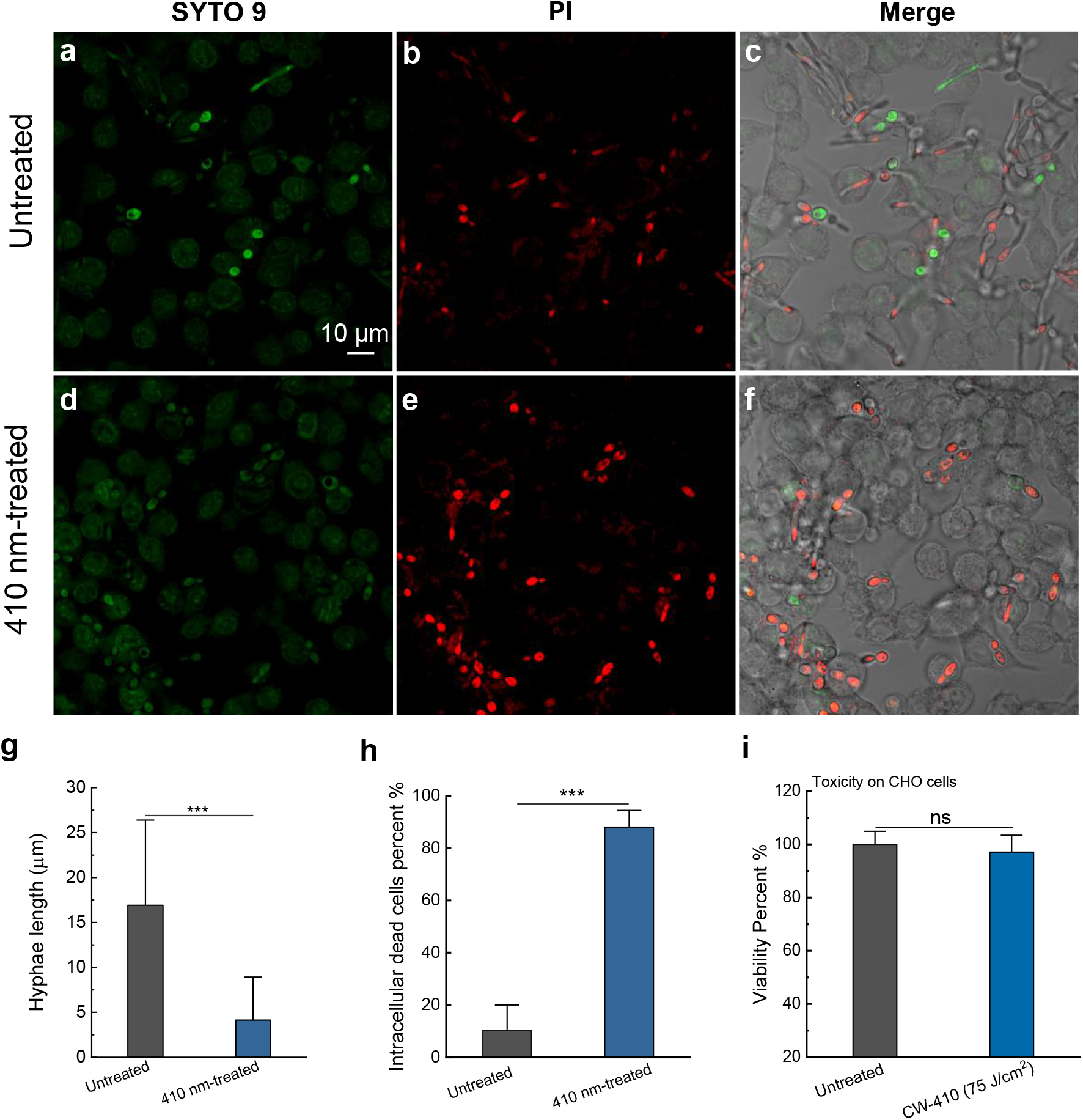
Elimination of intracellular *C. albicans* SC5314 by synergizing photoinactivation of catalase with host innate immune cells. **a-f**. Confocal imaging of live/dead *C. albicans* SC5314 after *C. albicans* SC5314 (**a-c**) infection of macrophages RAW 264.7 with or without 410 nm blue light pre-treatment for 1 hour at a multiplicity of infection of 10. **g**. Quantitative analysis of *Candida* hyphae length of the above two groups. **h**. Quantitative analysis of intracellular dead bacteria percent of the above two groups. **i**. CHO cell viability with or without 410 nm blue light treatment by MTT assay. Data: Mean+SD. N=3. Student unpaired *t*-test. *: *p*<0.05, **: *p*<0.01; ***: *p*<0.001.

It was also reported that catalase is an indispensable enzyme for *C. albicans* hyphal formation (47). Therefore, there are two plausible explanations for the photoinactivation of catalase-mediated macrophage killing. On one hand, photoinactivation of catalase might inhibit hyphal formation through catalase depletion, thus causing less macrophage rupture and lysis; one the other hand, ROS from macrophages can efficiently exert its antimicrobial effect against intracellular *C. albicans* without the protection of catalase. In short, photoinactivation of catalase holds the clinical potential for eliminating intracellular fungal cells.

### Photoinactivation of catalase reduces fungal burden in a *C. albicans*-induced mouse abrasion model

With these *in vitro* implications, we query the clinical utilization of photoinactivation of catalase against fungal infections. It was reported that *C. albicans*-caused infections usually start from superficial infections before developing severe systematic infections (48). Therefore, we chose a *C. albicans* infected mouse skin abrasion model (49) to evaluate our the treatment efficacy *in vivo*. Briefly, 10^6^ CFU of *C. albicans* SC5314 was applied to abraded mouse skin for three hours, after that different treatment schemes (120-J/cm^2^ 410 nm blue light, 0.5% H_2_O_2_, 410 nm blue light plus 0.5% H_2_O_2_) were then administered to the infected wounds (**Figure 7a**). Two hours after the second dose, mice were euthanized, and infected wounds homogenized and serially diluted onto *C. albicans*-specific BiGGY agar plates (50).

**Figure 7.**
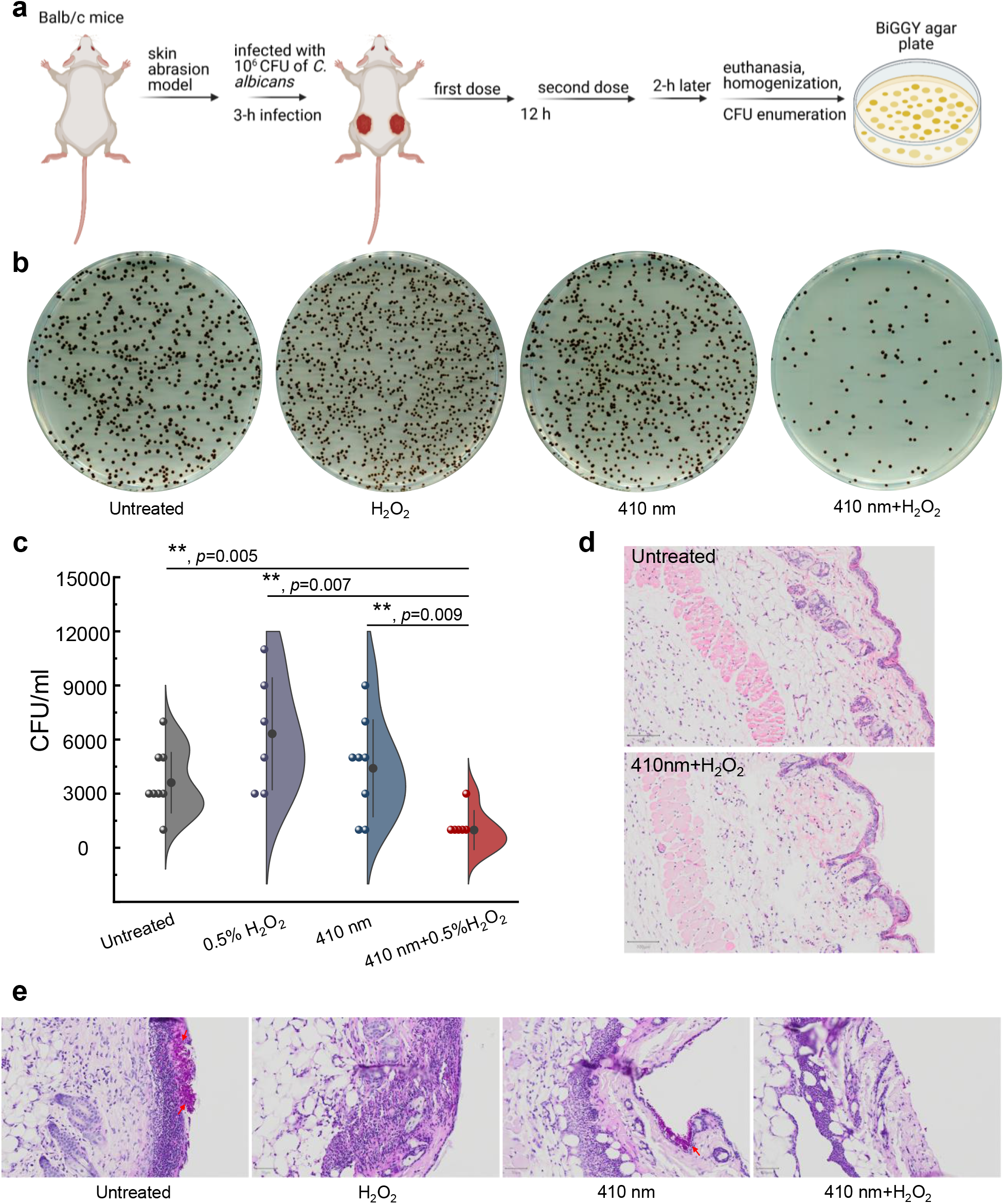
Photoinactivation of catalase and H_2_O_2_ synergistically reduces *C. albicans* burden in a mice skin abrasion model. **a**. Schematic illustration of development and subsequent treatment for *C. albicans*-induced mice skin abrasion. **b**. Spread BiGGY agar plates (*Candida* specific) from homogenized mice wounds after different treatment schemes. **c**. *C. albicans* CFU/ml of the homogenized mice wounds after different treatments in (**a**). **d**. H&E stained histology slides of the untreated mice skin and 410 nm blue light plus H_2_O_2_ treated mice skin. Scalar bar, 100 μm. **e**. H&E stained histology slides of *C. albicans* infected mice skins under different treatment schemes. Scalar bar, 50 μm. Fungal colonization area was pointed by red arrows. Data: Mean+SE from at least six replicates. Significant difference was determined through student unpaired *t*-test. *: *p*<0.05, **: *p*<0.01; ***: *p*<0.001. Outlier was determined through whisker box plot.

As shown in **Figure 7b**, *C. albicans* isolated from infected wounds were depicted as smooth black colonies on the BiGGY agar plates. Under the same dilution factor, 410 nm plus 0.5% H_2_O_2_ treated group apparently showed the lowest number of *C. albicans* colonies compared to the other treatment groups. Quantitative analysis of CFU/ml from homogenized wounds further confirmed that the fungal burden in the 410 nm blue light plus 0.5% H_2_O_2_ treated group was consistently the lowest compared to the other three groups (**Figure 7c**). These results suggest that photoinactivation of catalase is effective in reducing overall fungal burden in the clinical-relevant settings.

To interrogate whether the above treatment induces skin damage, we examined and further compared the normal mouse skin with or without 410 nm blue light (120 J/cm^2^) plus H_2_O_2_ treatment through hematoxylin and eosin (H&E) staining analysis. As shown in **Figure 7d**, epidermis, dermis, and subcutaneous tissues from the treated mouse skin appeared as intact as that of untreated one, indicating that there is no detectable toxicity from the 410 nm blue light (120 J/cm^2^) plus H_2_O_2_ (0.5%) treatment.

Meanwhile, we also employed H&E staining assay to visualize *C. albicans* infected mice skin from the above treatment groups. Yeast-form *C. albicans* turned into hyphal structures after seventeen hours of infection (**Figure 7e**), consistent with the fact that *C. albicans* usually transitions from yeast to hyphae form in disease settings (51). Of note, neutrophils or macrophages infiltration likely happened in the area adjacent to fungal infection site based on the histology analysis, further immunohistochemistry staining analysis is needed to understand which specific immune cells phagocytosed *C. albicans*. Strikingly, in the groups involved with 410 nm blue light treatment, hyphae-form *C. albicans* barely showed up. Only few *C. albicans* remained on the mice skin surface after 410 nm plus 0.5% H_2_O_2_ treatment. Collectively, photoinactivation of catalase reduces the virulence of *C. albicans*, thus rendering this pathogen highly susceptible to exogenous H_2_O_2_ attack in a clinically relevant mice fungal infection model.

### Consecutive blue light treatment does not induce resistance development by *C. albicans*

In most of clinical settings, multiple doses of treatments are usually necessary to ensure sufficient clearance of fungal cells. To understand whether continuous 410 nm treatment could induce fungal resistance, we examined the performance of *C. albicans* after six-consecutive-day blue light treatment (**Figure 8a**). First, CFU enumeration assay was performed on *C. albicans* at day 0, 3, 6 under different treatment schemes. Next, we compared the catalase amount of *C. albicans* at day 0 and day 6 through the Amplex Red catalase kit.

**Figure 8.**
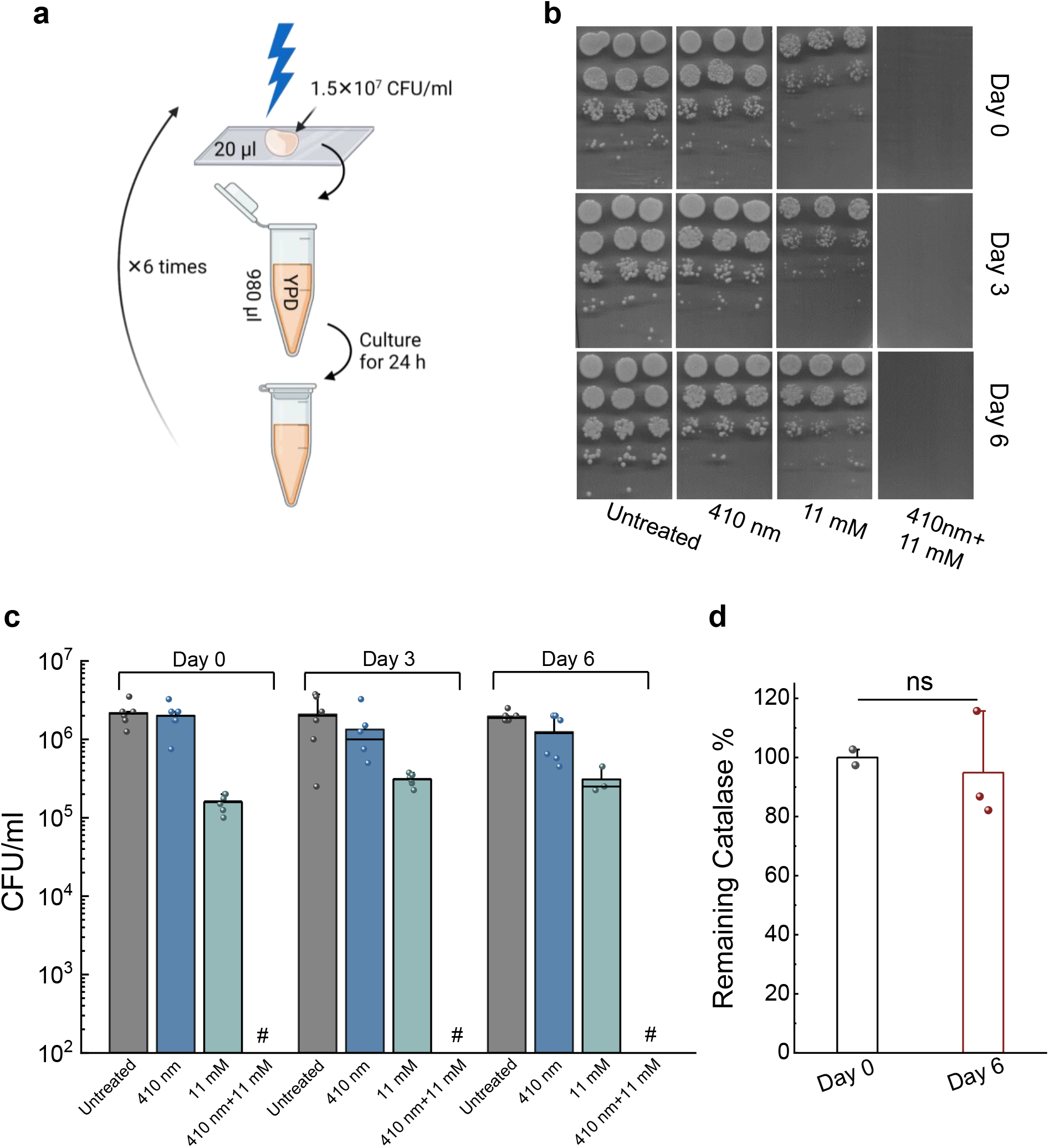
*Candida albicans* doesn’t develop resistance towards consecutive blue light treatments. **a**. Schematic illustration of the serial blue light treatment and passage experiment. **b**. Spread plates of CFU of *C. albicans* from Day 0, 3, 6 under different treatment schemes. **c**. Quantitative analysis of *C. albicans* CFU/ml under various treatments. Pound sign (#) indicates the CFU results are below the detection limit. **d**. Catalase amount comparison from *C. albicans* before and after six-consecutive blue light exposure. Student unpaired *t*-test. ns: not significant. 410 nm: 60 J/cm^2^.

As shown in **Figure 8b**, *C. albicans* from day 0, 3, 6 demonstrated similar sensitivity towards H_2_O_2_ treatment, and total eradication of *C. albicans* in the 410 nm blue light plus H_2_O_2_ treated group was achieved consistently. Quantitative analysis of CFU/ml results further consolidated the results, indicating that *C. albicans* doesn’t develop resistance towards consecutive blue light treatments.

To understand whether catalase amount has any difference before and after the consecutive treatments, we compared the catalase amount inside *C. albicans* at day 0 and day 6 using the Amplex Red catalase kit. Interestingly, we didn’t find significant difference between these two groups (**Figure 8c**), further suggesting that catalase, the molecular target of 410 nm blue light, is consistently expressed throughout the serial passage, and it is unlikely to develop resistance towards 410 nm blue light-based therapy.

## Discussion

Antifungal resistance has raised alarms and threats to clinicians and patients due to the limited arsenal of antifungal agents as well as current impeded antifungal development pipeline (52). For severely ill or immunocompromised patients, this situation especially worsens, and systematic candidiasis or candidemia might happen. Association of invasive aspergillosis accompanying COVID-19 patients as secondary infection have been reported (53). In addition, the global overuse of antifungals accelerates development of multidrug or pan-drug resistant fungal cells. Therefore, novel approaches to combat fungal infection are highly desired. Here, using wild-type *C. albicans* along with catalase-deficient *C. albicans*, we first identified catalase residing inside wide-ranging fungal species as a primary target of blue light. Its function in detoxifying intracellular H_2_O_2_ can be efficiently inactivated by blue light, especially around 410 nm range, which subsequently renders these fungal species highly susceptible to ROS-producing agents. In addition, the synergy between photoinactivation of catalase and H_2_O_2_ remains effective for various fungal species and strains, especially, the notorious clinical drug-resistant *C. auris* strains. Towards clinical applications, we found that photoinactivation of catalase boosts macrophage killing of intracellular *C. albicans*, and reduces *C. albicans* burden in a *C. albicans* infected mouse model of skin abrasion once combining with hydrogen peroxide. Consecutive blue light passage study indicates that neither catalase expression level nor cell susceptibility to blue light is changed for serial blue light treatment. These findings suggest the great potential of catalase-targeting blue light therapy as a new alternative to treat clinical multidrug resistant fungal infections.

Of note, blue light-mediated inactivation of *C. albicans* has been documented before (13). However, the underlying molecular target and the eradication mechanism for blue light are still unclear, which seriously hinders follow-up developments in improving its efficacy as well as its clinical translations. Based on our studies here we unveiled that blue light alone only exerts limited efficacy by inactivating catalase and ROS-producing agents are essential to be administered as adjuvants to synergize with 410 nm blue light for strikingly improved fungal eradication. For example, the light dose and settings we applied here are well below the ANSI standard (200 mW/cm^2^ for 410 nm), and blue light-mediated photoinactivation of catalase sensitizes *C. albicans* to non-fungicidal low-concentration H_2_O_2_ by around six orders of magnitude. As another example, AmB, the golden standard for antimycotic treatment for the most severe fungal infections (54), exhibits part of its killing efficacy through ROS burst after binding with ergosterol (55). However, high-dose AmB treatment often leads to acute renal failure as a well-documented serious complication (56). Our findings suggest an alternative approach to tune down the usage of AmB by photoinactivation of catalase. Moving further, photoinactivation of catalase also revives certain antifungal agents such as fluconazole against azole-resistant clinical *C. auris* strains (**Figure 5c-d**).

We found that multiple clinical *C. auris* strains are exceptionally sensitive to blue light when compared to the other fungal cells as total growth inhibition was obtained through PrestoBlue proliferation assay (**Figure 5**). However, it remains enigmatic regarding exactly why *C. auris* has such high sensitivity to blue light. Likely, it possesses relatively higher amount of catalase when compared to other *Candida* species. Nevertheless, further quantitative characterizations of catalase amount and related mRNA expression levels are helpful to address this query. *C. auris* also demonstrates a propensity to persist on skin or environmental surfaces, a unique feature to distinguish it from other *Candida* species, thus highly transmissible (57). Our data here demonstrates the high susceptibility of *C. auris* to blue light, and blue light alone can be developed as a novel approach for eradicating drug-resistant *C. auris* colonized on the skin or environmental surfaces.

We also found that photoinactivation of catalase boosts macrophage killing of intracellular *C. albicans*. Particularly, shortened hyphae length was observed inside the macrophages infected by the 410 nm pre-exposed *C. albicans*, suggesting the pivotal role of catalase plays in the transition from yeast-to hyphae-form *C. albicans*. It has been reported catalase gene disruption did not induce hyphae formation in yeast-form *Saccharomyces cerevisiae* (47). Therefore, further molecular-level study to further understand how 410 nm blue light treatment modulates such yeast-to-hyphae transformation can be pursued. In addition, significant number of yeast-form *C. albicans* were found dead inside macrophages, suggesting an effective 410 nm blue light-mediated macrophage phagocytosis. Other immune cells like neutrophils also produce high level of ROS when phagocytosing fungal cells (58). Therefore, we suspect that photoinactivation of catalase could assist neutrophils to eliminate intracellular fungal cells via a similar mechanism. Taken together, photoinactivation of catalase presumably enhances immune cells to phagocytose fungal cells, thus preventing the potential disseminated candidiasis.

In an independent and parallel study, we identified catalase in a wide range of bacterial species as a primary molecular target of blue light and further demonstrated the potential utilization of photoinactivation of catalase to eradicate various pathogenic bacteria through combination with H_2_O_2_-producing agents (59). The current work unveiled catalase as a primary target of blue light in wide-ranging fungal species, thus broadening the scope of utilization of photoinactivation of catalase to an even wider range of pathogenic microbes. Although catalase is identified as the primary target of blue light, we believe that other molecular targets other than catalase might still exist. This is evidenced by enhanced H_2_O_2_ killing of catalase-deficient *C. albicans* by 410 nm blue light. To identify these peripheral targets in the future, endogenous pigments could be further examined as intrinsic chromophores e.g. staphyloxanthin, have shown striking sensitivity to blue light-mediated photobleaching (60–63). It has also been reported that melanin is the major pigment inside *C. albicans* playing a myriad of biological functions (64). Photoinactivation or photomodulation of melanin might account for blue light-mediated killing as well. Another direction to pursue is to improve light penetration depth, as blue light currently can only penetrate a few hundred microns (65), thus limiting its applications only to superficial surfaces. For deep-seated fungal infections, pulsed blue laser with high-peak power might offer improved and deeper penetration compared to the continuous-wave light (61). Upconverting nanoparticles, capable of converting near-infrared red excitation into visible and ultraviolet emission (66), might suggest another potential way to treat deep-tissue fungal infections. In summary, our findings here suggest a fundamental molecular target and eradication mechanism of blue light elimination of multiple fungal species. These mechanistic insights create new possibilities to find highly effective treatment avenues for surface sterilization and for treatment of superficial fungal infections.

## Materials and Methods

### Blue light source

Continuous-wave (CW) blue light was delivered through a mounted 405 nm blue light LED (M405L4, Thorlabs) with an adjustable collimation adapter (SM2F32-A, Thorlabs) focusing the illumination region to a ~1 cm^2^ region. A T-Cube LED driver (LEDD1B, Thorlabs) allowed for adjustable light fluencies up to 500 mW/cm^2^.

### Fungal strains and chemicals

#### Fungal strains

*Candida albicans* SC5314 (wild type, American Type Culture Collection (ATCC)). Catalase-deficient *Candida albicans* 2089 (△*cat1*) are from Dr. Alistair Brown lab at University of Exeter. All the other strains are from Dr. Michael K. Mansour’s lab (clinical strains from Massachusetts General Hospital (MGH), Boston, MA) including all the *Candida auris* strains: *Candida auris 1*_MGH, *Candida auris 1* (AR-0381), *Candida auris 2* (AR-0382), *Candida auris 3* (AR-0383), *Candida auris 4* (AR-0384), *Candida auris 5* (AR-0385), *Candida auris 6* (AR-0386), *Candida auris 7* (AR-0387), *Candida auris 8* (AR-0388), *Candida auris 9* (AR-0389), *Candida auris 10* (AR-0390). *Candida albicans C15, Candida albicans C16, Candida albicans C17. C. glabrata* (ATCC 2001), *Candida glabrata C1, Candida glabrata* C2, *C. tropicalis* (H3222861), *C. parapsilosis* (F825987), *C. lusitaniae* (S1591976), *C. lusitaniae* (AR-0398), *C. haemulonii* (AR-0393), *C. haemulonii* (AR-0395), *Candida duobushaemulonii* (AR-0394), *C. krusei* (AR-0397). All cell lines used in this study, including the RAW 264.7 murine macrophages, Chinese hamster ovary (CHO) cells were purchased directly from the ATCC.

#### Chemicals

Amphotericin B (A9528-100MG, Sigma Aldrich). Hydrogen peroxide (CVS Health). DMSO (D8418-500ML, Sigma Aldrich). YPD broth (Y1375, Sigma Aldrich). YPD Agar (Y1500, Sigma Aldrich). Phosphate buffered saline (BP399500, Fisher Scientific). 10% formalin (HT501128-4L, Sigma Aldrich). Sodium chloride solution (S8776, Sigma Aldrich). Miconazole nitrate salt (M3512, Sigma Aldrich). Fluconazole (86386, Acros Organics). PrestoBlue reagent (A13261, ThermoFisher). 3-Amino-1,2,4-triazole (3-AT, A8056, Sigma Aldrich). Bovine liver catalase (C1345, Sigma Aldrich). LIVE/DEAD cell viability kit (L7007, ThermoFisher). BiGGY agar (73608, Sigma Aldrich). Amplex Red Catalase Assay (A22180, Thermo Fisher Scientific). DMEM (2186822, Gibco). HI FBS (2273356P, Gibco). HEPES (54457, Sigma Aldrich).

#### Fungal culture

*C. albicans* was routinely streaked at 30 °C onto the yeast peptone dextrose (YPD) agar and sub-cultured in YPD broth overnight to stationary phase. Stationary-phase *Candida* strains were cultured into mid-log phase prior use by 1:20 dilution by YPD broth. Fungal cell density was adjusted based on the optical density at 660 nm (OD_600_). The suspension was centrifuged, washed with phosphate-buffered saline (PBS), and re-suspended in PBS at the cell density of 10^6^ ~10^7^ CFU/ml before proceeding all the treatments.

#### Quantitation of remaining catalase percentage

Measurement of remaining catalase percentage was primarily quantified through the use of an Amplex Red Catalase Assay (A22180, Thermo Fisher Scientific) according to its standard protocol. Briefly, solutions containing catalase (either bovine liver catalase or catalase-positive fungal culture with 1×10^6^ cell density) were treated with blue light, after which 25 μl of the light treated solution was incubated with 40 μM of H_2_O_2_ for two hours at 30°C in dark inside a 96-well plate. Following H_2_O_2_ treatment, 50 μl of a reaction stock containing 100 μM of Amplex Red and 0.4 U/mL of horseradish peroxidase were added to each well and then incubated for 30 minutes at 37°C. Following incubation, the fluorescence of each well was measured at an excitation wavelength of 545 nm and an emission wavelength of 590 nm. A negative control containing the 1×reaction buffer (0.1 M Tris-HCl) and an untreated positive control (normal fungal cells without blue light exposure) also treated with the assay in order to determine the remaining catalase percentage. Remaining catalase percentage was calculated through the following equation:

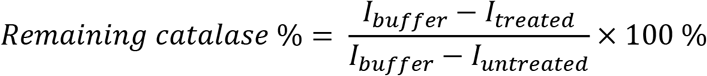

### Imaging of intracellular H_2_O_2_

*C. albicans* with and without 410 nm exposure (30 J/cm^2^) were incubated with H_2_O_2_ (22 mM, 30-min incubation). After that, *C. albicans* were stained with an intracellular H_2_O_2_ kit (MAK164-1KT) for 30 min. Fungal cells were then washed and sandwiched between a cover slide and polylysine-coated cover glass. Confocal laser scanning microscopy was then applied to image intracellular H_2_O_2_.

#### Bubbling test, CFU enumeration assay, short-term incubation assay, Checkerboard broth dilution assay and time killing assay to validate the mechanism of catalase inactivation in catalase inhibited or knocked-out Candida strains

*Candida albicans* SC5314 and *Candida albicans △cat1* were cultured overnight in YPD broth at 30°C using an orbital shaking incubator (250 rpm).

For the **bubbling test**, after overnight culturing, both the wild type *C. albicans* SC5314 and *Candida albicans △cat1* were washed by PBS and adjust the cell density to 10^7^ /ml, placed 1 ml of fungal solution into cuvettes with 80 μl of 3% H_2_O_2_, and then incubated at room temperature for 30 min before observation.

For the **CFU enumeration**, the next day after overnight culture, fungal cultures were resuspended 1:20 in YPD for another 5~6 hours and cultured into mid-log phase. Then fungal cells were washed, suspended in PBS and adjusted at the optical density by 660 nm (OD_660_) to ~10^8^ CFU/ml, after which a 10 μl of aliquot was placed on a glass cover slide and then treated with CW-410 nm (100 mW/cm^2^ 5min). Following light treatment, the droplet was collected to a tube containing PBS or PBS supplemented with H_2_O_2_. H_2_O_2_ ranged from 0.69 mM to 22mM according to the sensitivity of fungal strains. Samples were then incubated for 30 minutes at 30°C in the orbital shaking incubator, after which the treated and untreated fungal samples were 10-fold serially diluted inside a 96-well plate, plated on YPD agar plates in a 30°C static incubator for 24~36 hours prior to enumeration. All the CFU results were read at the same time for each patch of experiment.

#### Short-term incubation assay

In order to determine the impact of short term H_2_O_2_ exposure on *C. auris* following light induced catalase inactivation, *C. auris* 2 was incubated overnight in YPD broth. The next day 200 μl of *C. auris* was centrifuged and resuspended in 1 ml of 1×PBS. A 10 μl aliquot of *C. auris* suspension was placed on a glass coverslip and exposed to 30 J/cm^2^ of CW-410 (200 mW/cm^2^, 2.5 min). Following exposure, the aliquot was transferred to 990 μl of PBS and vortexed. This process was repeated for the non-light treated *C. auris* as well. After the creation of the light treated and untreated *C. auris* stock, 300 μl of stock was separated and treated with 0.1% H_2_O_2_ for 1 min, after which the H_2_O_2_ treated suspension was serially diluted and plated to determine changes in overall CFU following exposure.

For the **catalase inhibition** assay, 3-Amino-1,2,4-triazole (3-AT) was applied to both untreated and H_2_O_2_ containing groups at the concentration 50 mM), and cultured with *C. albicans* SC5314 for 4 hours in a 30°C shaking incubator. After treatment, the treated suspensions were spun down and resuspended in fresh PBS, after which a 10 μl aliquot of untreated *C. albicans* was treated with 30 J/cm^2^ of CW-410 (100 mW/cm^2^, 5 min) and then diluted in 990 μl of PBS. Alongside the light treated dilution, an untreated dilution and 3-AT treated dilution were also produced. These dilutions were treated with 44 mM of H_2_O_2_ for 2 h at 30°C, after which the samples were serially diluted and plated to quantify the overall CFU enumeration.

For the **time killing assay**, *Candida albicans* SC5314 and *Candida albicans △cat1* were cultured into mid-exponential phase in YPD broth at 30°C using an orbital shaking incubator (250 rpm). Then the fungal culture was washed and adjusted based on OD_660_ to ~10^8^ CFU/ml, after which a 10 μL aliquot was placed on a glass cover slide and then treated with CW-410 nm (100 mW/cm^2^, 5 min). Following light treatment, the droplet was collected to a tube containing PBS or PBS containing 11mM H_2_O_2_. Samples were shaken and incubated at 30°C for up to 4 hours. During the incubation, 60 μL aliquots were removed from each sample tube for serial dilution and CFU plating at the 0 min, 20 min, 40 min, 1 h, 1.5 h, 2 h and 4 h time points.

For the **checkerboard assay**, *Candida albicans* SC5314 and *Candida albicans △cat1* were cultured into mid-exponential phase in YPD broth. Then fungal culture was washed and adjusted based on OD_660_ to ~10^7^ CFU/ml as stock solution and stored on ice. An aliquot of 5 μl fungal stock solution was exposed with 410 nm light at a power of 500 mW/cm^2^ for 0 min, 2.5 min, 5 min, 10 min, 20 min, 40 min, then collected with 995 μl of YPD broth to a final cell density of ~5×10^4^/ml and plated into a 96-well plate in row. Then a two-fold serial dilution was performed from 22 mM of H_2_O_2_ to the following concentrations: 11 mM, 5.5 mM, 2.8 mM, 1.4 mM, 0.7 mM, 0.4 mM, 0.2 mM, 0 mM. Time-course OD_660_ was recorded by the plate reader over 18 hours. The combinational behavior between blue light and H_2_O_2_ from both wild type *C. albicans* and catalase mutant was then calculated based on the following equation:

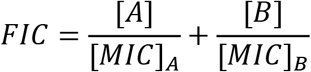

Where the FIC stands for fractional inhibitory concentration, indicating the synergy level of two treatments. (FIC < 0.5: Synergy; FIC > 4: Antagonism; 0.5 < FIC < 4: Additive or indifference)

### Time-course PrestoBlue proliferation assay of log phased and stationary phased Candida species

Clinical fungal strains from MGH Dr. Michael K. Mansour’s lab were used in this part to test the broad band of our phototherapy, and a high throughput resazurin-based PrestoBlue assay was applied. Both stationary- and exponential phase fungal cells are included as well. Stationary phase *Candida* species were prepared by overnight culturing at 30°C using an orbital shaking incubator (250 rpm); and the log phase fungal cells were prepared the next day by 1:20 in YPD for another 5~6 hours and cultured into mid-log phase, some of the *C. auris* strains took longer time to grow. All cells were collected and washed twice, then the cell density was adjusted to ~10^8^/ml prior to blue light exposure. A 10 μl of aliquot was placed onto a glass cover slide and then treated with CW-410 nm (100 mW/cm^2^, 5min) or not. Following light treatment, the droplet was collected to a tube containing 990 μl of PBS to reach the fungal cell density of ~10^6^/ml. H_2_O_2_ ranging from 11 mM to 44 mM was added to the wells in the first row and a two-fold dilution was conducted, the initial concentrations were chosen according to the sensitivity of fungi strains. Samples were then incubated for 4 hours at 30°C, after which an aliquot of 90 μl of YPD and 10 μl of PrestoBlue reagent were added to each well, followed by a time-course recording of the fluorescence signal (Excitation: 590 nm) for 24 h. Calibration growth curves with different initial cell density was also conducted in order to calculate the original CFU of all the *Candida* strains.

### Intracellular fungi assay

To assess the fungal intracellular killing with murine macrophage RAW 264.7 cell line, *C. albicans* SC5314 was collected from an exponentially growing culture and washed with PBS. Macrophages were cultured in Dulbecco’s Modified Eagle medium (DMEM) alongside 10% fetal bovine serum (FBS) until 90% confluence was achieved. Macrophages were pre-washed with serum-free DMEM media immediately before infection, and infected by *C. albicans* SC5314 (MOI=10) with and without 410 nm treatment (35 mW/cm^2^, 8min). Then co-culture was incubated at 37°C in a humidified incubator to allow for the phagocytosis of fungal cells. After two hours, the infection co-culture was removed and replaced with normal growth media (DMEM supplemented with 10% FBS, 10 mM HEPES). A Live (SYTO 9)/Dead (PI) staining assay was to utilize to visualize the live and dead fungal cells inside macrophages, respectively. Briefly, after fixation with 10% formalin following infections, samples were permeabilized with 0.1% Triton-X for 3 min at room temperature. After that, a Live/Dead fluorescence kit (Thermo Fisher Scientific, L7007) was utilized to stain the intracellular fungal cells and confocal laser scanning microscope (FV3000, Olympus) was employed to visualize the stained samples.

#### Mammalian cell toxicity assay

To evaluate the potential toxicity of 410 nm exposure and short term, high concentration H_2_O_2_ exposure against mammalian cells, an MTT assay was performed using Chinese Hamster Ovary (CHO) cell line. CHO cells were cultured in Dulbecco’s Modified Eagle medium (DMEM) and 10% fetal bovine serum (FBS) until a high confluence was obtained. CHO cells were then removed via trypsin, quantified, and diluted in serum-free DMEM media to a cell concentration of 1×10^6^ cells/ml. In a treated 96-well plate, 100 μl of cell suspension was added to each well, providing 1×10^5^ cells per well. The cells were allowed to adhere overnight at 37°C with 5% CO_2_.

The next day, the media was removed from each well and the cells were washed twice with PBS to minimize potential reactions between the phenol red present in DMEM and the blue light treatment. Washing conditions performed on one set of cells were also performed on all other sets to maintain consistency. Following washing, all wells were filled with PBS and the light treated wells were then exposed to 30 J/cm^2^ of CW-410 (200 mW/cm^2^, 2.5 min) After light treatment, PBS was removed and replaced with DMEM. For the H_2_O_2_ treated groups, 0.1% H_2_O_2_ in DMEM was added to the H_2_O_2_ treatment groups for 1 min, after which the media was immediately removed and the wells were washed three times with PBS to minimize potential remaining H_2_O_2_ remaining. Three-fold washing was also applied to the other treatment groups. After washing, the wells were filled with 100 μl of fresh DMEM and the cells were allowed to recover from treatment over the course of 24 hours at 37°C with 5% CO_2_. Once 24 hours have passed, an MTT viability assay was performed, where the DMEM was replaced with fresh DMEM alongside 0.5 mg/ml of MTT (Life Technologies, M6494) and incubated at 37°C for 4 h. Once upon completion, the DMEM/MTT solution was removed and 100 μl of filtered DMSO was mixed in and provided with 45 min to dissolve the crystalized formazan, after which absorbance measurements were quantified via plate reader at 590 nm. The assay was performed in replicates of N=4.

#### *In vivo* murine infection model and histology

Animal studies were approved by the Institutional Animal Care and Use Committee (IACUC) at Boston University and were performed in the Animal Science Center (ASC) in the Charles River Campus (CRC). The abrasion fungal infection model was used. Twenty BALB/c mice (Jackson Laboratories, 000651) were shaved on the dorsal side and placed under anesthesia, then a sterile #15 sterile scalpel was used to generate two ~1×1 cm^2^ abrasion wounds on both left and right by carefully scraping the epidermis of the skin until reddish dots appeared without drawing blood out of skin barrier. Once wounds were generated, a 10 μl aliquot containing 2×10^8^/ml of exponential phase *C. albicans* SC5314 in PBS was placed onto the abrasion wound, and spread evenly across the wound with a pipette tip and air dried. Then 20 mice were divided into four treatment groups, each consisting of 5 mice (N=10 wounds in total): Untreated, 410 nm treated, H_2_O_2_ treated, and 410 nm plus H_2_O_2_ treated. Light treatment was applied by positioning mice under 200 mW/cm^2^ (ANSI standard for human skin) 410 nm LEDs for 10 min (total 120J/cm^2^, 5-min exposure and 5-min break and then 5-min exposure again in order to prevent potential photodamage). For H_2_O_2_ treatment, 10 μl of 0.5% H_2_O_2_ was evenly distributed on the infected wounds and air dried. Combinational treatment consisted of the application of previously described blue light treatment (120 J/cm^2^) followed immediately by the administration of H_2_O_2_. Treatments were applied to mice twice, the first treatment being applied 4 hours following infection, and the second treatment being applied 20 hours after the infection. 2 hours after the second treatment, mice were euthanized and wound tissue was harvested, either homogenized and serially diluted for CFU studies or fixation by 10% neutral buffered formalin for histology studies. CFU enumeration was performed on *C. albicans*-specific BiGGY agar plates (73608, Sigma Aldrich).

The potential phototoxicity of the treatment on the skin was evaluated by hematoxylin and eosin (H&E) histology assay. Same assay was applied as well to examine the location of fungal cells in the infection site. The untreated mouse skin received the same two treatments as the wound site, and during tissue harvesting, this region was excised and preserved in 10% buffered formalin. Formalin fixed samples for both untreated and treated normal skin tissues (N=6) and fungi infected skin tissues were submitted to the Boston Medical Center (BMC) for histology processing, periodic acid-Schiff (PAS) staining for fungal infected skin and hematoxylin and eosin (H&E) staining for normal skin. Histology slides were then visualized by Olympics VS120 slide scanner provided by Micro/Nano Imaging (MNI) lab in Biomedical Engineering Core Facilities under in Boston University, and then the images were visualized and analyzed by a pathologist Dr. Ivy Rosales at MGH.

### Statistical analysis

Statistical analysis was conducted through student unpaired *t*-test and One-way ANOVA. *** means significantly different with the *p*-value < 0.001. ** means significantly different with the *p*-value < 0.01. * means significantly different with the *p*-value < 0.05. ns means no significance.

## Acknowledgments

This work was partly supported by RO1AI141439 to J.-X.C, and RO1AI132638 to M.K.M. We kindly acknowledge Dr. Alistair Brown lab at University of Exeter for providing the catalase-deficient *C. albcians* strains. Research reported in this publication was also supported by the Boston University Micro and Nano Imaging Facility and the office of the director, National Institute of Health, National Institute of Health under Award Number S10OD024993.

## Author contributions

P.-T.D. and J.-X.C. conceived the synergistic therapeutic treatment between photoinactivation of catalase and H_2_O_2_ or certain antifungals. M.K.M. provided all the clinical fungal strains including the Candida auris strains and experimental discussions. P.-T.D. and J.H. discovered that catalase from catalase-positive fungi could be ubiquitously inactivated by 410 nm. P.-T.D. characterized catalase photoinactivation and intracellular fungi assay. P.-T.D. and Y.W.Z. conducted the mechanism study by the catalase-deficient *C. albicans* strains. P.-T.D. and S.J. conducted the 3-AT study. Z.D., N.A., P.-T.D., and Y.Z. prepared the clinical fungal samples together. P.-T.D., Y.W.Z. and J.H. conducted all the *in vitro* PrestoBlue assay. P.-T.D., Y.W.Z., and S.J. did data analysis of Prestoblue assay. Y.W.Z., P.-T.D. and S.J. conducted the *in vivo* mice abrasion experiments and histology assay. Y.W.Z. conducted the histology slides scanning and preliminary analysis. P.-T.D. and J.-X.C. co-wrote the manuscript. G.L. provided constructive suggestions over the project and manuscript. All authors read and commented on the manuscript.

## Supplementary information

**Supplementary Figure 1.**
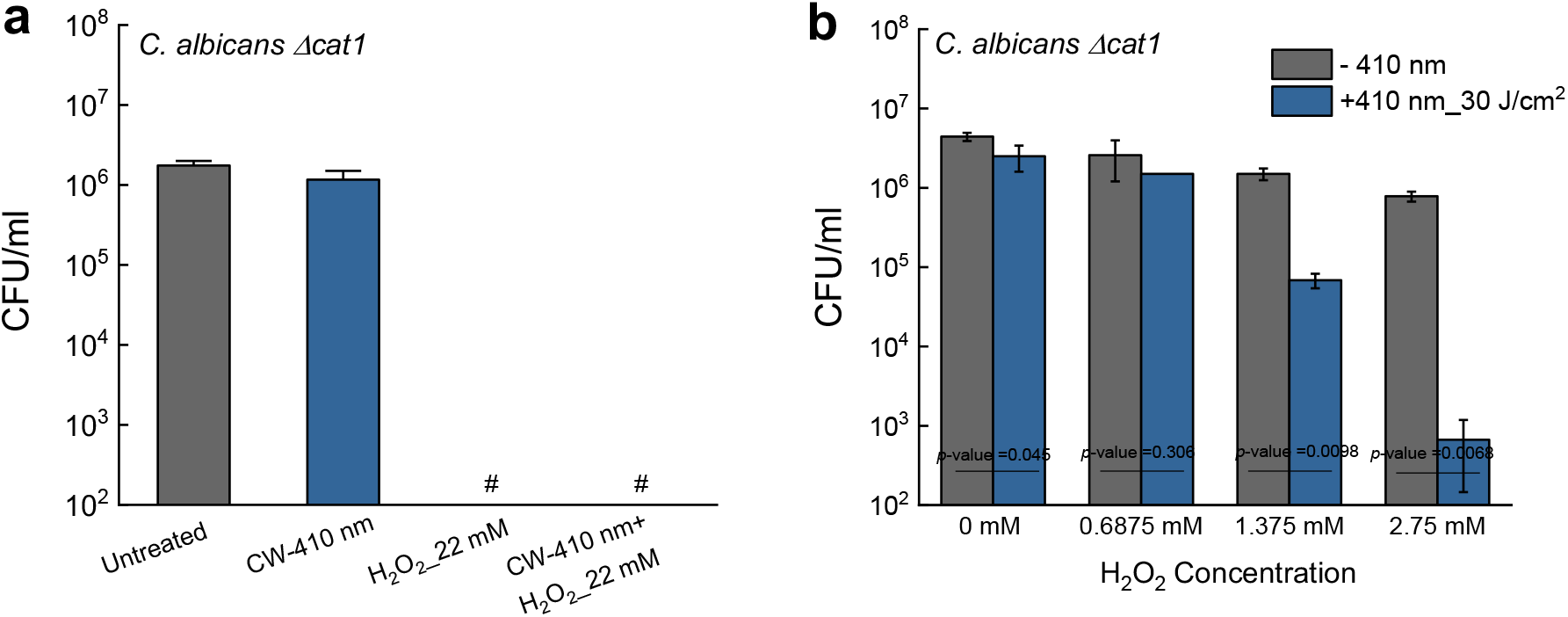
CFU/ml of catalase-deficient *C. albicans Δcat1* under different treatment schemes. Data: Mean+SD. N=3. Student unpaired *t*-test. *: *p*<0.05, **: *p*<0.01; ***: *p*<0.001.

**Supplementary Figure 2.**
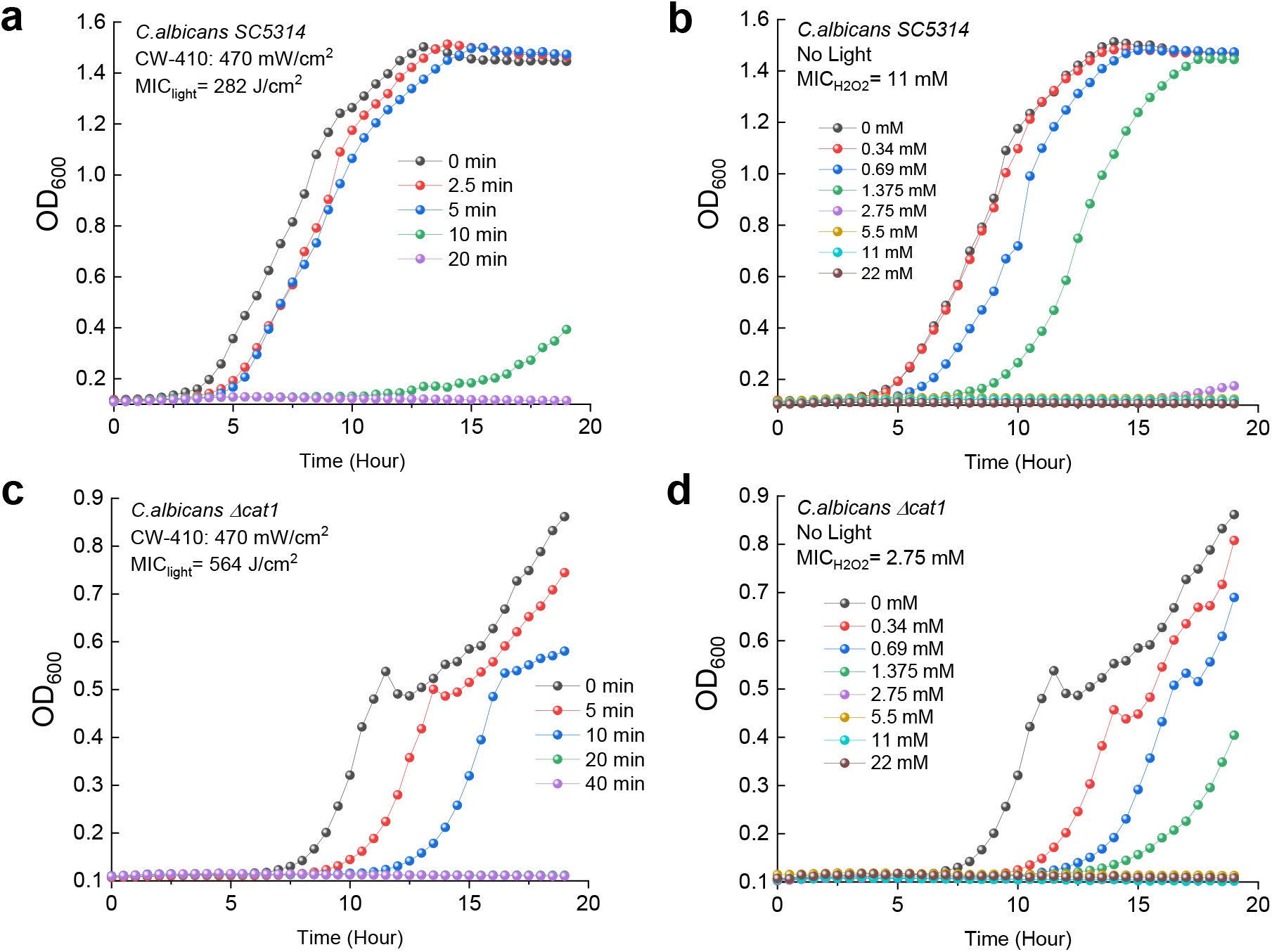
Minimal inhibitory concentrations of 410 nm exposure and H_2_O_2_ against *C. albicans* SC5314 and catalase-deficient *C. albicans Δcat1*.

**Supplementary Figure 3.**
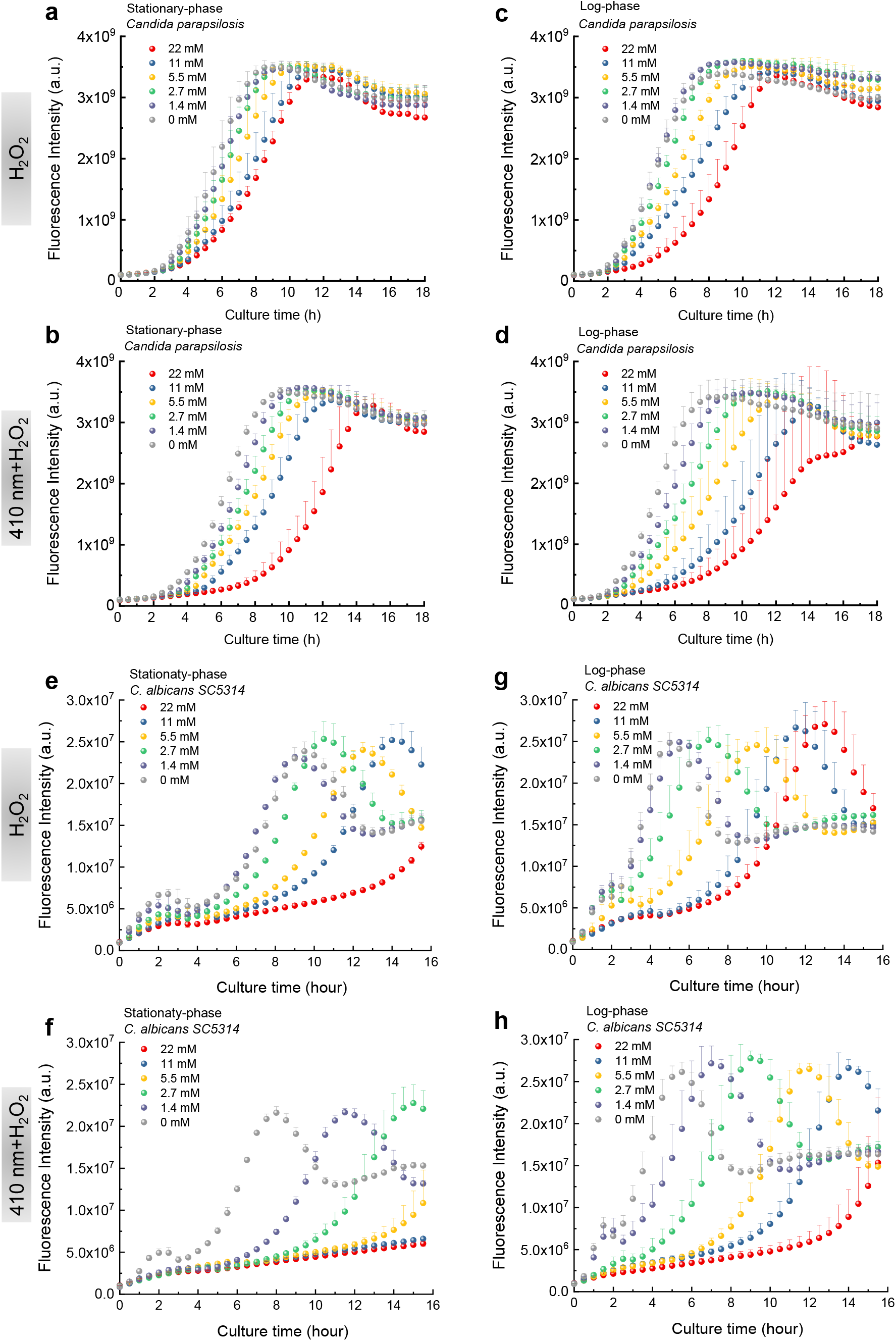
Time-course PrestoBlue fluorescence intensity from various fungal species under the treatment of 410 nm and H_2_O_2_ of various concentrations.

**Supplementary Figure 4.**
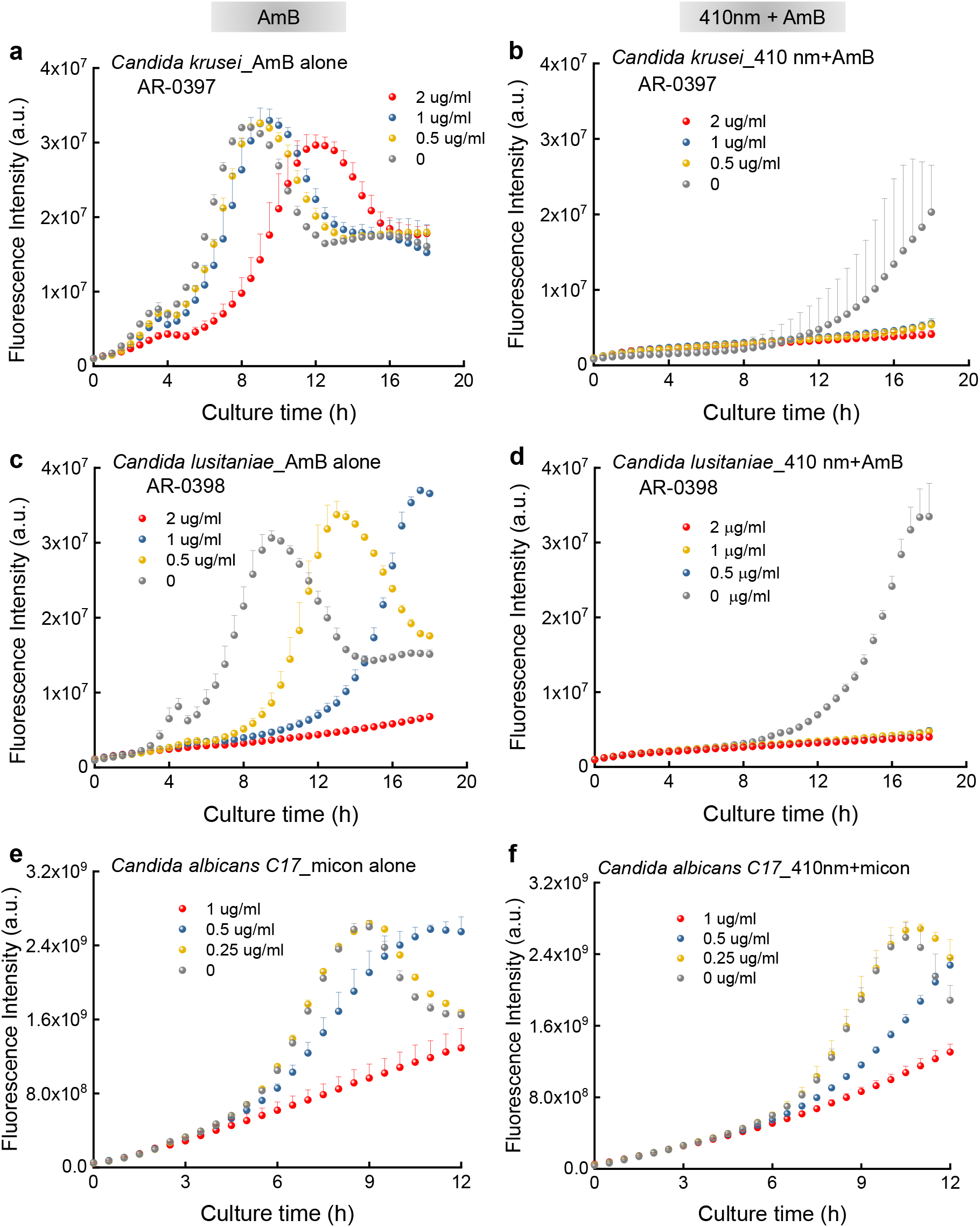
Time-course PrestoBlue fluorescence intensity from various fungal species under the treatment of 410 nm and antifungal drugs.

**Supplementary Figure 5.**
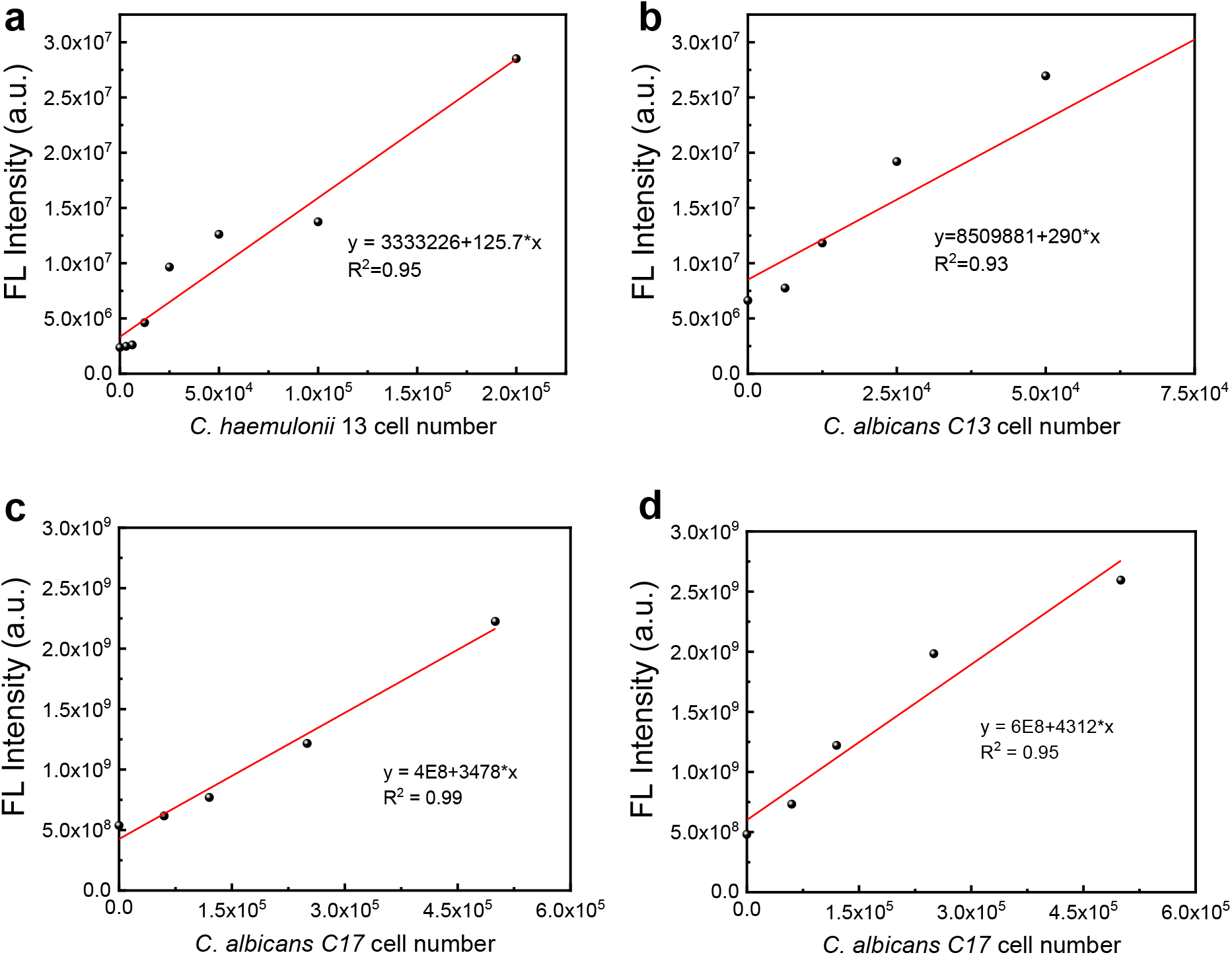
Calibration curves of *Candida spp*. CFU/ml versus PrestoBlue fluorescence intensity.

**Supplementary Figure 6.**
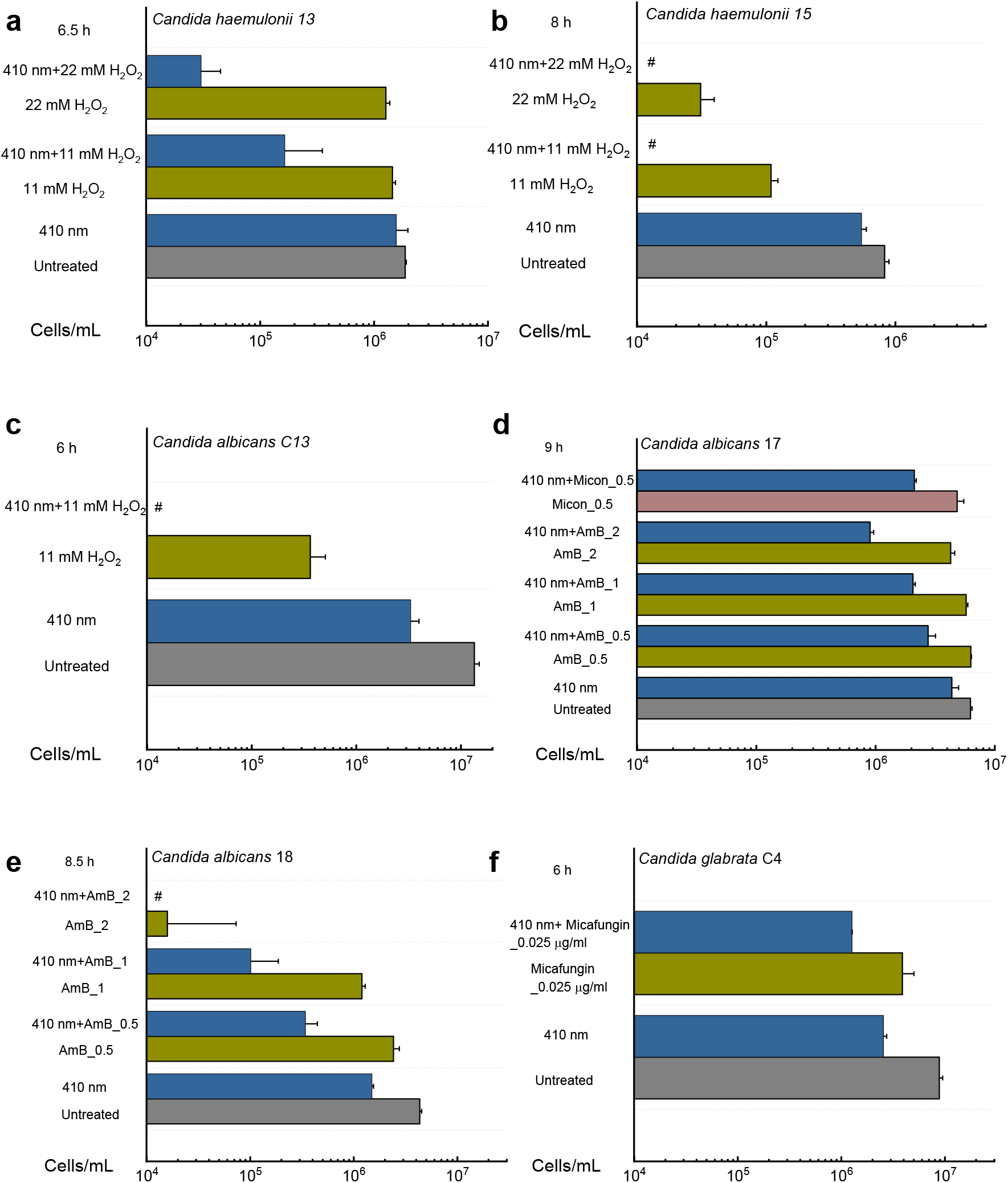
Derived Candida cells/mL after different treatments based on the calibration curves. Data: Mean+SD. N=3. Pound sign (#) means below the detection limit.

**Supplementary Figure 7.**
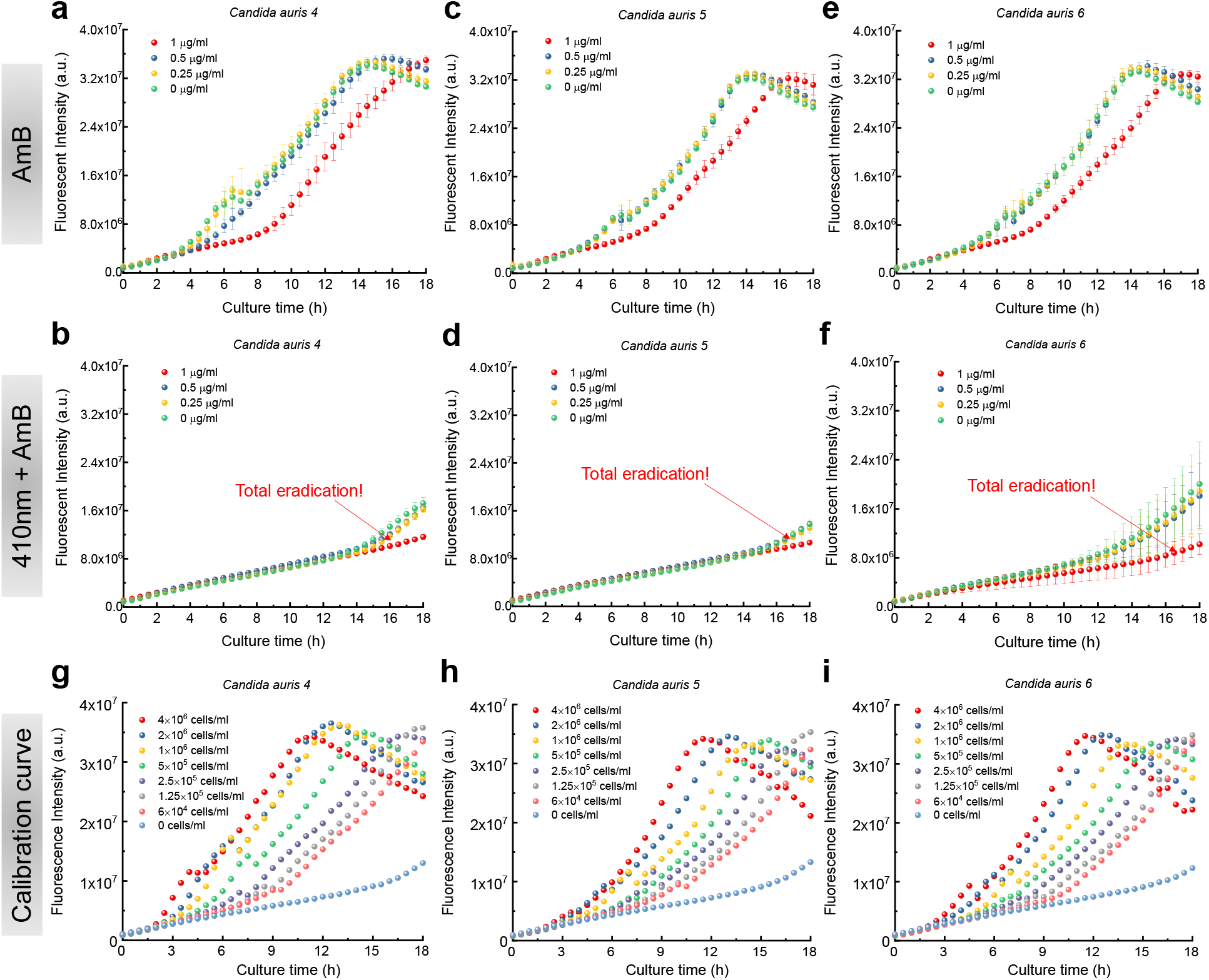
Time-course PrestoBlue fluorescence intensity from various *Candida auris* strains under the treatment of 410 nm and antifungal drugs.

**Supplementary Figure 8.**
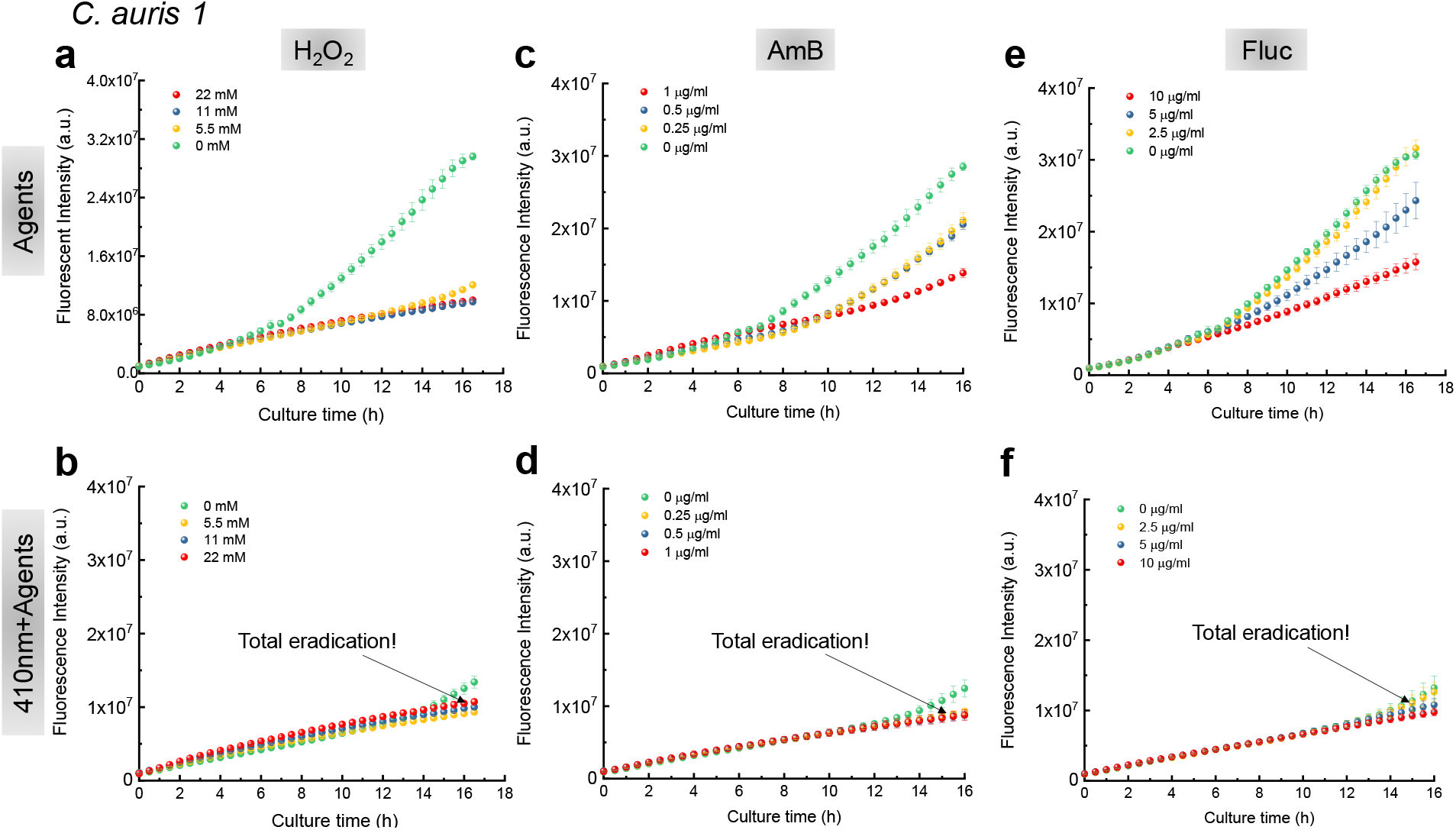
Time-course PrestoBlue fluorescence intensity from *Candida auris* 1 under the treatment of 410 nm and antifungal agents.

**Supplementary Figure 9.**
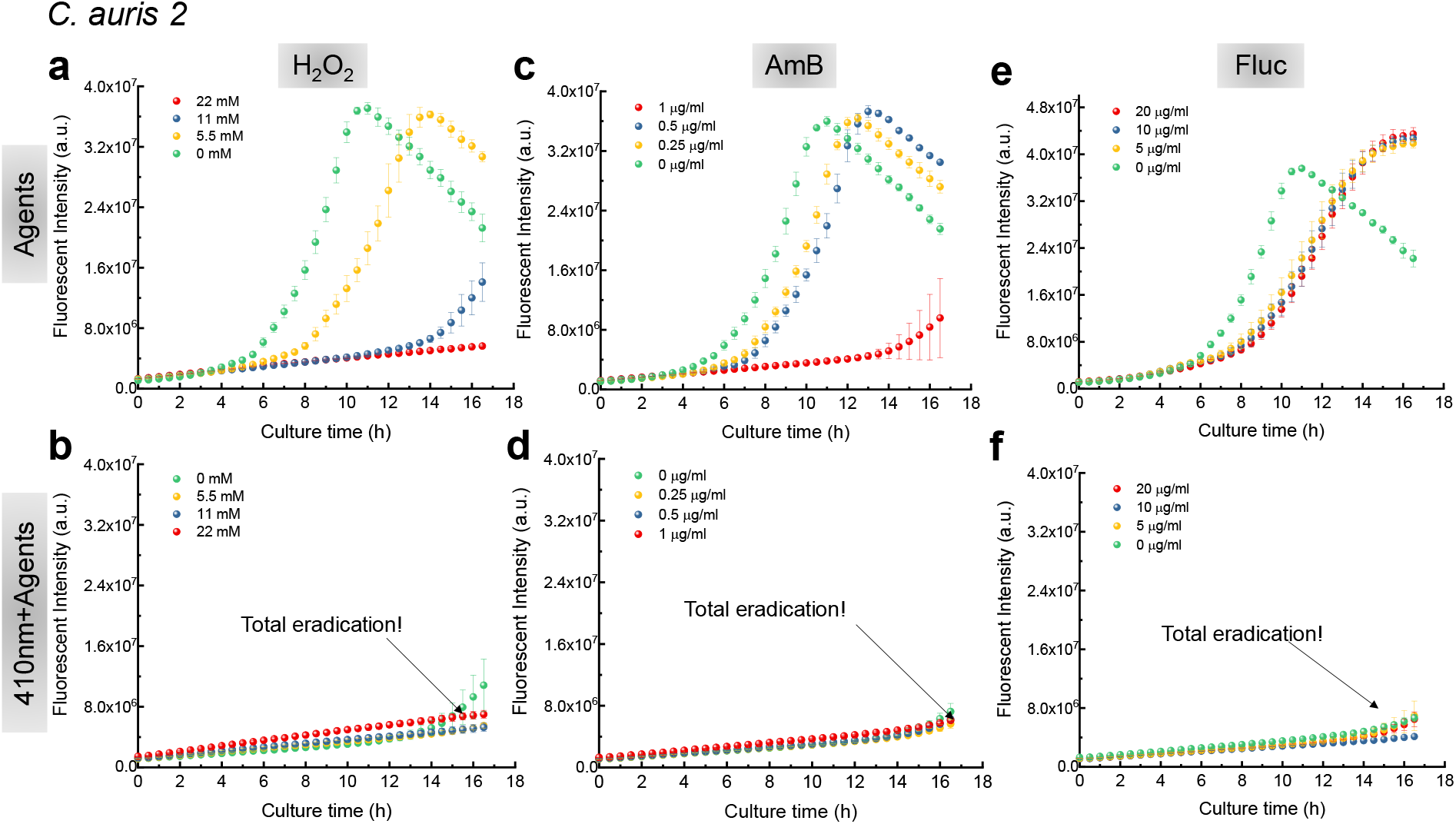
Time-course PrestoBlue fluorescence intensity from *Candida auris 2* under the treatment of 410 nm and antifungal agents.

**Supplementary Figure 10.**
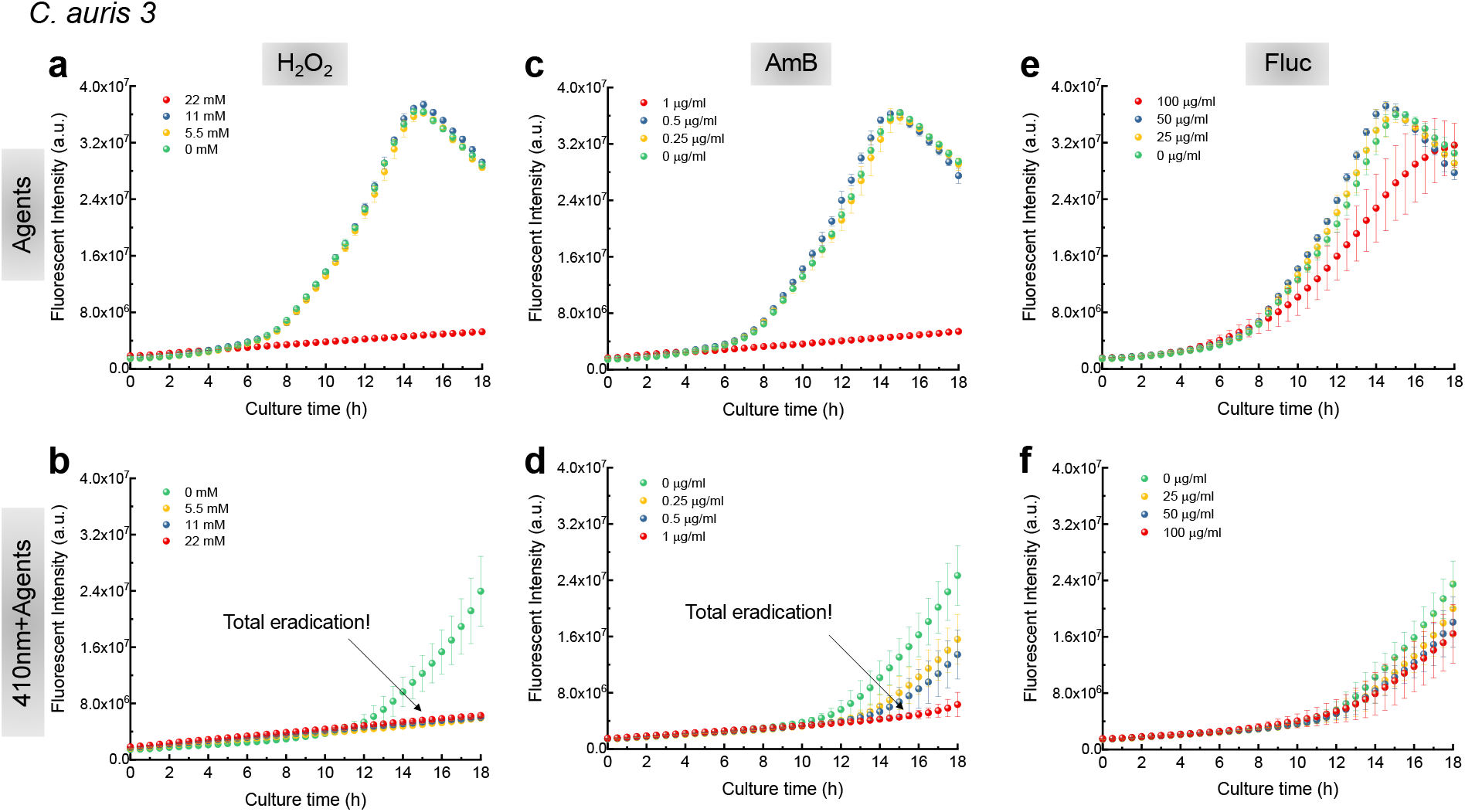
Time-course PrestoBlue fluorescence intensity from *Candida auris* 3 under the treatment of 410 nm and antifungal agents.

**Supplementary Figure 11.**
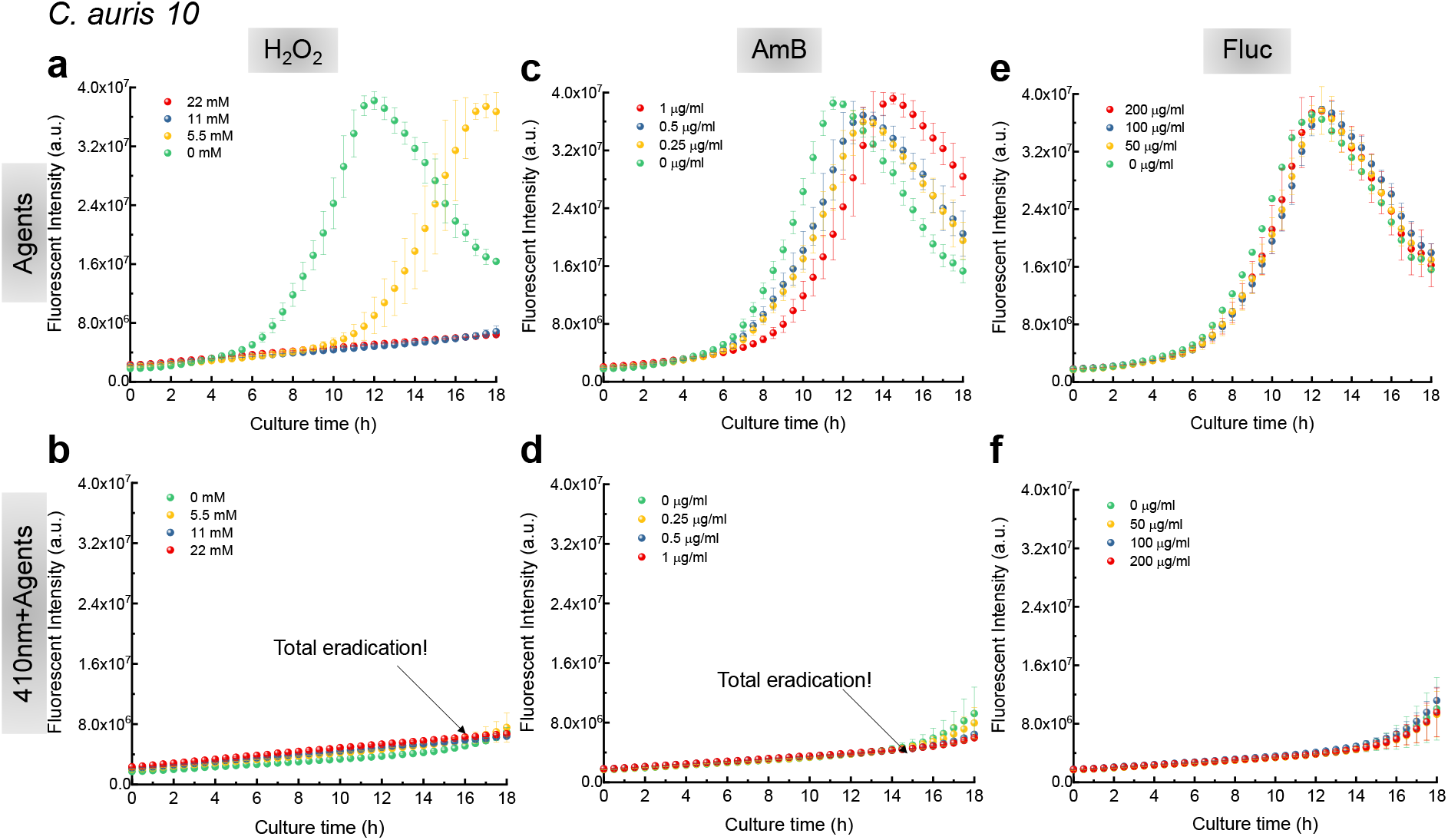
Time-course PrestoBlue fluorescence intensity from *Candida auris* 10 under the treatment of 410 nm and antifungal agents.

**Supplementary Figure 12.**
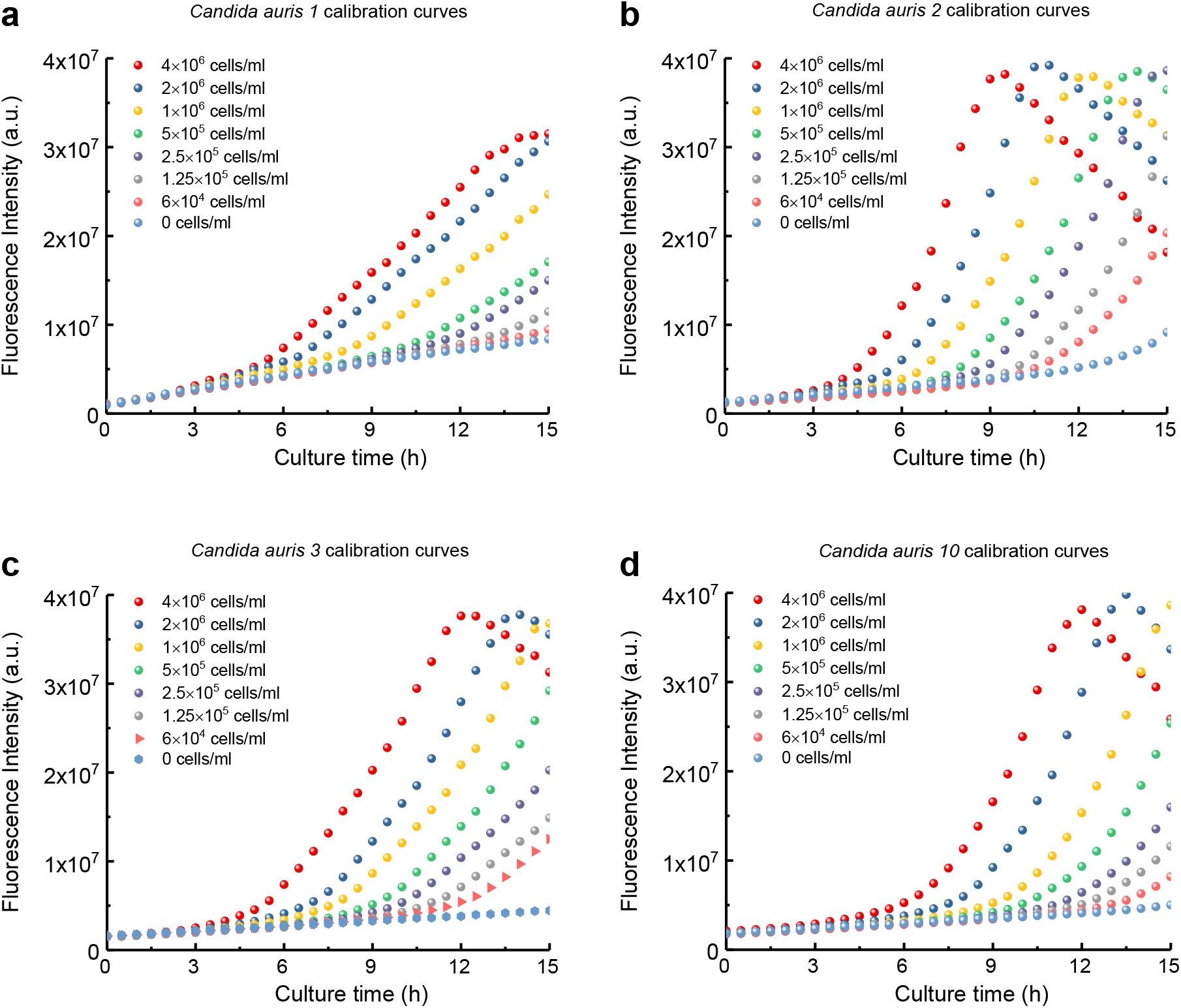
Time-course PrestoBlue fluorescence intensity of *Candida auris* strains at specific concentrations.

**Supplementary Figure 13.**
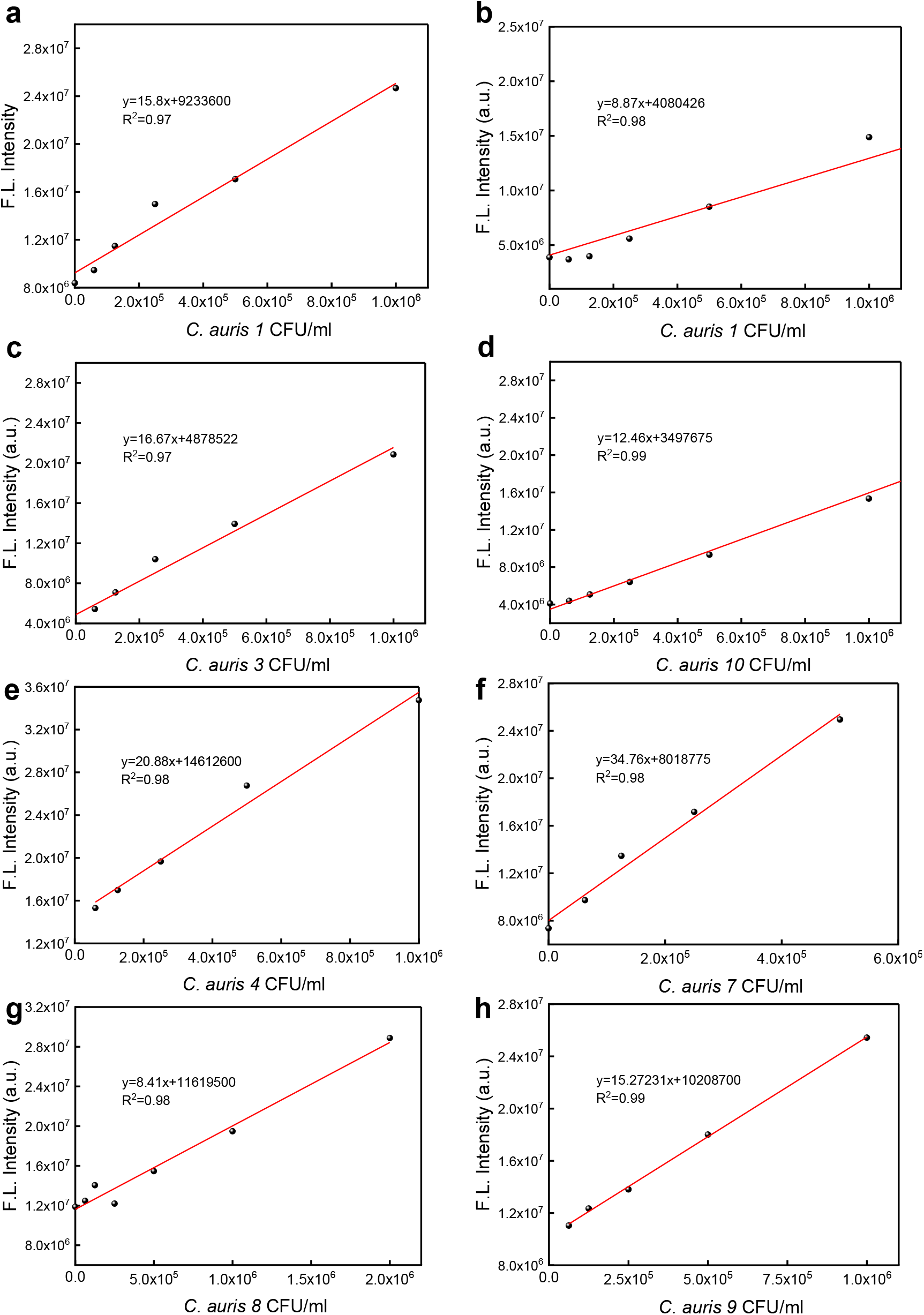
Calibration curves of *C. auris* CFU/ml versus prestoblue fluorescence intensity.

